# Disruption of lysosomal proteolysis in astrocytes facilitates midbrain proteostasis failure in an early-onset PD model

**DOI:** 10.1101/2022.08.26.505472

**Authors:** Gustavo Morrone Parfitt, Elena Coccia, Camille Goldman, Kristen Whitney, Ricardo Reyes, Lily Sarrafha, Ki Hong Nam, Soha Sohail, Drew Jones, John F Crary, Alban Ordureau, Joel Blanchard, Tim D Ahfeldt

## Abstract

Accumulation of advanced glycation end products (AGEs) on biopolymers accompany cellular aging and drives poorly understood disease processes. Here, we studied how AGEs contribute to development of early on-set Parkinson’s Disease (PD) caused by loss-of-function of DJ1, a protein deglycase. In induced pluripotent stem cell (iPSC)-derived midbrain organoid models deficient for DJ1 activity, we find that lysosomal proteolysis is impaired, causing AGEs to accumulate, α-synuclein (α-syn) phosphorylation to increase, and proteins to aggregate. These processes are at least partly driven by astrocytes, as DJ1 loss reduces their capacity to provide metabolic support and triggers acquisition of a pro-inflammatory phenotype. Consistently, in co-cultures, we find that DJ1-expressing astrocytes are able to reverse the proteolysis deficits of DJ1 knockout midbrain neurons. In conclusion, astrocytes’ capacity to clear toxic damaged proteins is critical to preserve neuronal function and their dysfunction contributes to the neurodegeneration observed in PD.

## INTRODUCTION

Aging is the strongest risk factor for developing neurodegenerative diseases such as Alzheimer’s Disease (AD) and Parkinson’s Disease (PD) (Reeve et al., 2014). Accordingly, investigating how biological mechanisms of aging are interconnected with the progression of neurodegenerative diseases is an active research area with significant therapeutic potential. Recently, glycation, a process in which aldehyde metabolites and nucleophiles become attached to biopolymers through non-enzymatic reactions, has come into focus as a disease-driving mechanism. The accumulation of such advanced glycation end products (AGEs) on nucleotides, lipids, and proteins is known to damage cell function and is a feature of normal aging (Chaudhuri et al., 2018). However, aberrant AGEs accumulation has also been linked to multiple human pathologies including aging-related neurodegenerative diseases such as PD. Both genetic and environmental factors contribute to the development of PD (Billingsley et al., 2018), presenting challenges for identifying disease mechanisms that can be targeted therapeutically (Schneider et al., 2020). Recently, however, mutations in the α-synuclein gene (*SNCA*), the first identified causal mutations in PD (Billingsley et al., 2018), were reported to impair proteome maintenance and cause protein aggregation (Stojkovska et al., 2021), highlighting protein quality control and autophagy as convergent disease-driving pathways in sporadic PD.

The protein DJ1, encoded by the *PARK7* gene, is causally linked to development of early-onset PD by loss-of-function (LOF) mutations (Bonifati et al., 2003; Lockhart et al., 2004), but how reduced function of DJ1 contributes to PD pathogenesis is not understood. Because a conserved cysteine residue in DJ1 is known to be frequently oxidized, it was initially suggested that DJ1 can sense the oxidative state of the cell (Bandopadhyay et al., 2004; Canet-Avilé et al., 2004). Indeed, highly oxidized sulfonated (−SO_3_^−^) forms of DJ1 are associated with its inactivation and are increased in the cortex of PD patients when compared to age-matched controls (Bandopadhyay et al., 2004; Zhou et al., 2006). More recently, several studies have established that DJ1 has glyoxalase activity and may function as a deglycating enzyme that protects DNA, proteins, or lipids from harmful glycation, although some controversy remains about DJ1 substrates and activity levels (Andreeva et al., 2019; Hasim et al., 2014; Jun and Kool, 2020; Richarme et al., 2015, 2017; Sharma et al., 2019). In addition, studies using human-induced pluripotent stem cell (iPSC) models demonstrate that DJ1 impacts on oxidative stress pathways in neurons, particularly dopamine oxidation and autophagy pathways (Ahfeldt et al., 2020; Burbulla et al., 2017). Collectively, these findings suggest that decreased DJ1 activity resulting from (−SO_3_^−^) oxidation may contribute to PD pathogenesis.

Cell death in PD is restricted to discrete neuronal populations in a few brain areas, and more specifically to dopaminergic neurons (DNs) in the substantia nigra (SN), which has led to a neuro-centric view of PD pathology (Surmeier et al., 2017). However, in recent years, genome-wide association studies have increasingly implicated glial-associated genes in PD (Nalls, et al., 2011). These findings are consistent with post-mortem analysis of PD cases, which often identifies astrocytes with a mild increase of GFAP in the SN, and accumulation of α-syn and PACRG in intracellular inclusions, unique features compared to the abundant astrogliosis observed in other neurodegenerative diseases, (Song et al., 2009b; Tong et al., 2015; Hishikawa et al., 2001). More recently, a scRNA-seq analysis identified a population of CD44/S100A6-high reactive astrocytes in the midbrain of PD patients (Smajić et al., 2022). In addition, dysregulation of glial neurometabolic coupling and neuro-immune interactions were reported to have a crucial role in PD initiation and the death of the most vulnerable dopaminergic neurons (di Domenico et al., 2019; Kam et al., 2020; Streubel-Gallasch et al., 2021; Tsunemi et al., 2020; Wilson et al., 2019). Together, these observations suggest that astrocytes have a causal role in PD pathogenesis.

Although DJ1 is abundantly expressed in the CNS, it’s most highly expressed in astrocytes of the cortex and SN (Bandopadhyay et al., 2004). In animal models, DJ1 activity in astrocytes is protective against chemical-induced lesions to DNs of the SN (Choi et al., 2018; De Miranda et al., 2018). Conversely, DJ1 knockout (KO) in mouse astrocytes leads to an exacerbated inflammatory response and impaired lesion repair (Choi et al., 2019). Thus, although DJ1 has a well-established role as a protective protein in response to increased oxidative stress levels, more recently, its glyoxalase activity and role as a deglycating enzyme has been described. These findings propose that glycation stress is the core mechanism that underlies disease initiation and progression in DJ1 loss of function (LOF) early-onset PD.

The development of midbrain human brain organoids (hMIDOs) enabled the recapitulation of human midbrain tissue features such as, mature TH-positive neurons and neuromelanin accumulation (Ahfeldt et al., 2020; Jo et al., 2016). hMIDOs recapitulate PD-related phenotypes in various PD mutations in long-term cultures such as, reduction in TH^+^ dopaminergic and accumulation in α-syn oligomers and phosphorylated forms(Ahfeldt et al., 2020; Mohamed et al., 2021). Together with the scalability and stability of organoid models make an ideal model for long-term studies and drug screening for PD pathology.

Here, we leverage a comprehensive panel of assays to investigate metabolic function in patient-derived human brain tissue and DJ1 LOF organoid iPSC models. We show that protein quality control pathways in astrocytes are defective in PD-associated. Accordingly, when DJ1 is missing or mutated, AGE accumulation and a-syn aggregation ultimately cause DN death. Our findings unravel the pathogenic role of astrocytes in PD and uncover potential therapeutic strategies.

## RESULTS

### DJ1 KO human midbrain organoids have reduced autophagy and PD-associated α-syn phenotypes

To study how DJ1 LOF impacts on cellular processes in the absence of potential contributions from PD-associated genetics, we generated homozygous and heterozygous DJ1 knockout iPSC lines via CRISPR-mediated genome editing of the BJSIPS iPSC line (originally derived from a healthy non-PD male). We confirmed the DJ1 KO via Sanger sequencing and Western blotting (Figures S1A-B). In addition, we obtained the healthy male donor-derived hiPSC line KOLF 2.1J (Pantazis et al., 2021) and the isogenic line harbouring the PD-associated DJ1 L166P LOF point mutation. We selected two clones which we hereafter refer to as L166P-1 and L166P-2.

To model effects of DJ1 LOF on midbrain PD pathology, we generated human midbrain organoids (hMIDOs) using an established midbrain patterning protocol (Ahfeldt et al., 2020; Sarrafha et al., 2021) (Figure 1A). Consistent with previous work, hMIDOs expressed midbrain markers FOXA2, LMX1A, the mature midbrain marker NURR1, and generated TH^+^ neurons. At day 40, we found no differences in expression of FOXA2, LMX1A, and NURR1 in DJ1 KO hMIDOs compared to control organoids (Figure S1C). At day 100, we confirmed the presence of mature dopaminergic neurons by measuring dopamine levels by mass spectrometry, although levels did not differ between control and KO organoids (Figure S1D). In contrast, we detected fewer NURR1-positive cells in organoids generated from the two DJ1 L166P clones (L166P-1 and L166P-2) (Figure S1C). By day 100, astrocytes emerged, as reflected by GFAP expression, which became strongly increased by day 200 (Figure 1B).

**Figure 1.**
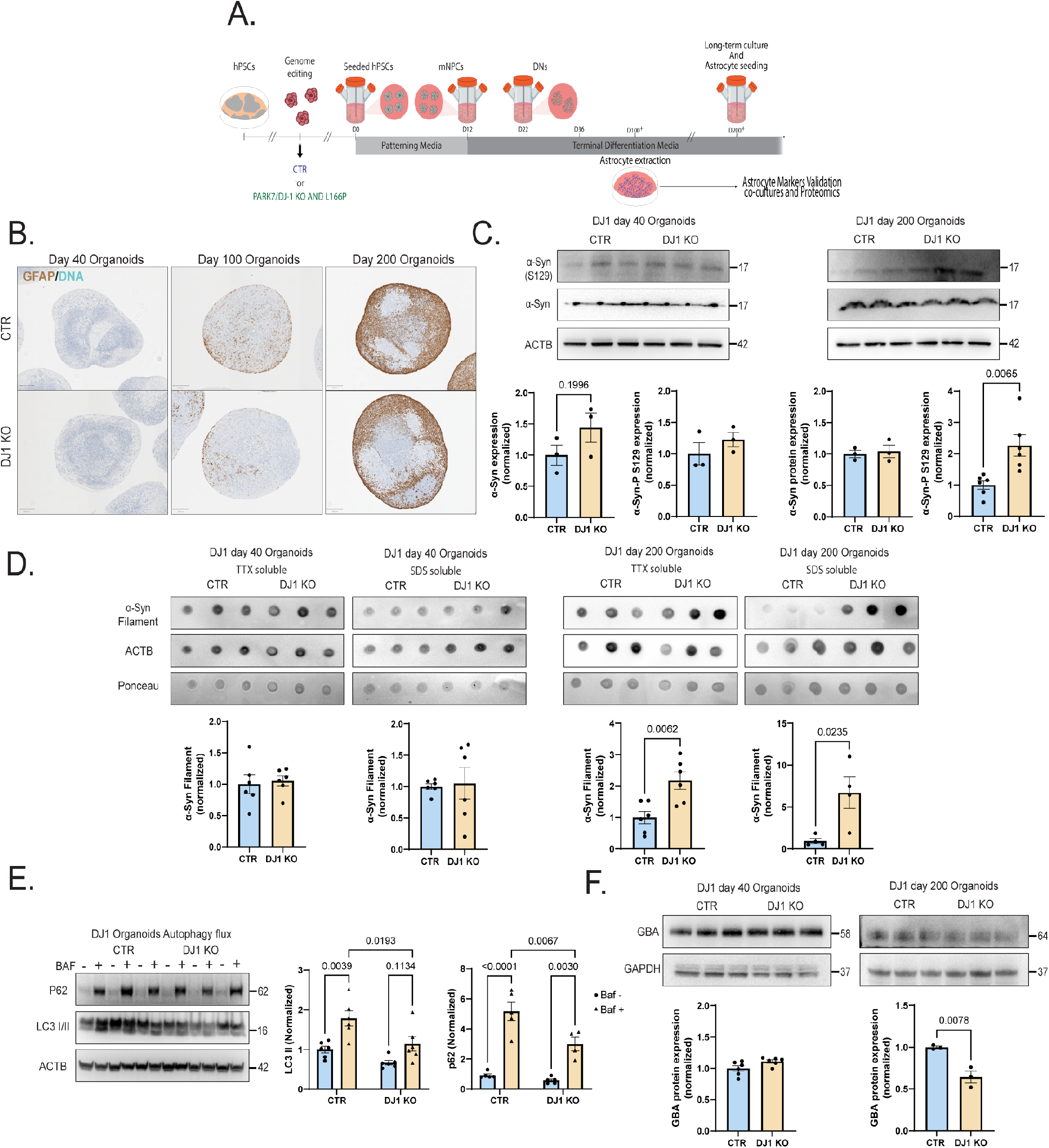
A. Micrography GFAP staining of CTR and DJ1 KO day 40, 100, and 200 midbrain organoids. B. Midbrain differentiation and astrocytes extraction protocol schematic. C. Immunoblots for α-syn, phospho-α-syn (S129), and actin (ACTB) loading control for CTR and DJ1 KO day 40 and day 200 midbrain organoids. C. Dot blots for oligomeric α-syn and actin (ACTB)/Ponceau loading control for CTR and DJ1 KO day 40 and day 200 midbrain organoids from TTX soluble and SDS soluble fractions. E. Immunoblots for P62, LC3 I/II, actin (ACTB) loading control in BAF − and + treated CTR and DJ1 KO day 100 midbrain organoids. F. Immunoblots for GBA and GAPDH loading control for CTR and DJ1 KO day 40 and day 200 midbrain organoids. All data are represented in mean ± S.E.M, data points are individual well differentiation, and the p-value was reported on the graph highlighted comparison.

The aggregation of α-synuclein protein in the midbrain is a hallmark of PD (Lashuel et al., 2013). Consistent with this phenotype, we found that aged DJ1 KO hMIDOs contained significantly increased levels of monomeric phosphorylated α-syn at day 200 relative to isogenic control midbrain organoids (Figure 1C). In contrast, we observed no significant difference in α-syn phosphorylation at day 40, suggesting that day 40 and day 200 midbrain organoids represent early and late disease stages (Figure 1C). Given that phosphorylation of α-synuclein at the S129 residue is correlated with α-syn turnover and accumulation in PD patients’ brain (Oueslati, 2016), we next assessed α-synuclein aggregation via western blotting. In day 40 organoids, we did not observe any significant difference in α-synuclein between DJ1 KO and controls (Figure 1D). However, in TTX- and SDS-soluble fractions of day 200 DJ1 KO midbrain organoids, levels of aggregated α-syn had increased (Figure 1D).

Lysosome pathways are essential to multiple forms of autophagy and known to process α-syn monomers and aggregates. Consistently, we previously detected lysosomal function as a dysregulated pathway using proteomics analysis of human embryonic stem cells (hESCs) and day 35 hESC-derived hMIDOs lacking DJ1 (Ahfeldt et al., 2020). Here, we re-analyzed our proteomics dataset using pathway analysis to identify a wider range of cellular processes potentially related to lysosomal dysfunction (Figure S1E). To investigate whether accumulation of α-synuclein aggregates in DJ1 KO hMIDOs was due to alterations in autophagy, we treated day 100 hMIDOs with bafilomycin A1 (BAF), which blocks lysosomal activity by inhibiting lysosome acidification. We then quantified levels of the autophagosome marker LC3 II, which is generated through lipidation of LC3 I and essential for autophagosome formation. In BAF-treated samples, our analysis shows that LC3 II levels are lower in DJ1 KO relative to CTR hMIDOs, indicative of an impaired autophagosome maturation (Figure 1E). Similarly, accumulation of the autophagy substrate P62 was decreased in BAF-treated KO hMIDOs (Figure 1E). Additionally, we observed depletion of the mature form of the lysosomal enzyme GBA in day 200 hMIDOs but no alterations at day 40 (Figure 1F), consistent with earlier reports in stem-cell derived neuronal PD models (Mazzulli et al., 2011; Stojkovska et al., 2022). Taken together, a decrease in LC3 lipidation, P62 accumulation, and depletion of mature GBA all point to a suppressed autophagy flux in DJ1 KO hMIDOs which would hinder degradation of protein aggregates.

### DJ1 KO midbrain organoids accumulate protein glycation damage

Oxidative damage, which compromises both proteome stability and cell viability (Dahl et al., 2015; Fleming et al., 2022), has been proposed to drive neurodegeneration through accumulation of advanced glycation end-products (AGEs) (Takeuchi and Yamagishi, 2008; Vicente Miranda et al., 2017). In PD patients, AGEs and their corresponding RAGE receptors accumulate in the SN and cortex (Dalfó et al., 2005), potentially due to DJ1 dysfunction, given its reported role as a deglycase that maintains AGE homeostasis (Richarme et al., 2017). Here, we quantified Methylglyoxal-derived hydroimidazolone (MGH) levels, an initial advanced glycation modification, in iPSCs and human midbrain organoids via dot blots. Although MGH levels were equivalent in DJ1 KO and control in both iPSCs and day 40 midbrain organoids, MGH protein glycation accumulated significantly in day 100 and day 200 DJ1 KO hMIDOs relative to controls (Figure 2A). In addition, low molecular weight (< 60 KDa) MGH bands were increased specifically in day 200 DJ1 KO organoids (Figure 2B) demonstrating that DJ1 loss of function result in AGE accumulation and concomitant synuclein pathology in late-stage organoids.

**Figure 2.**
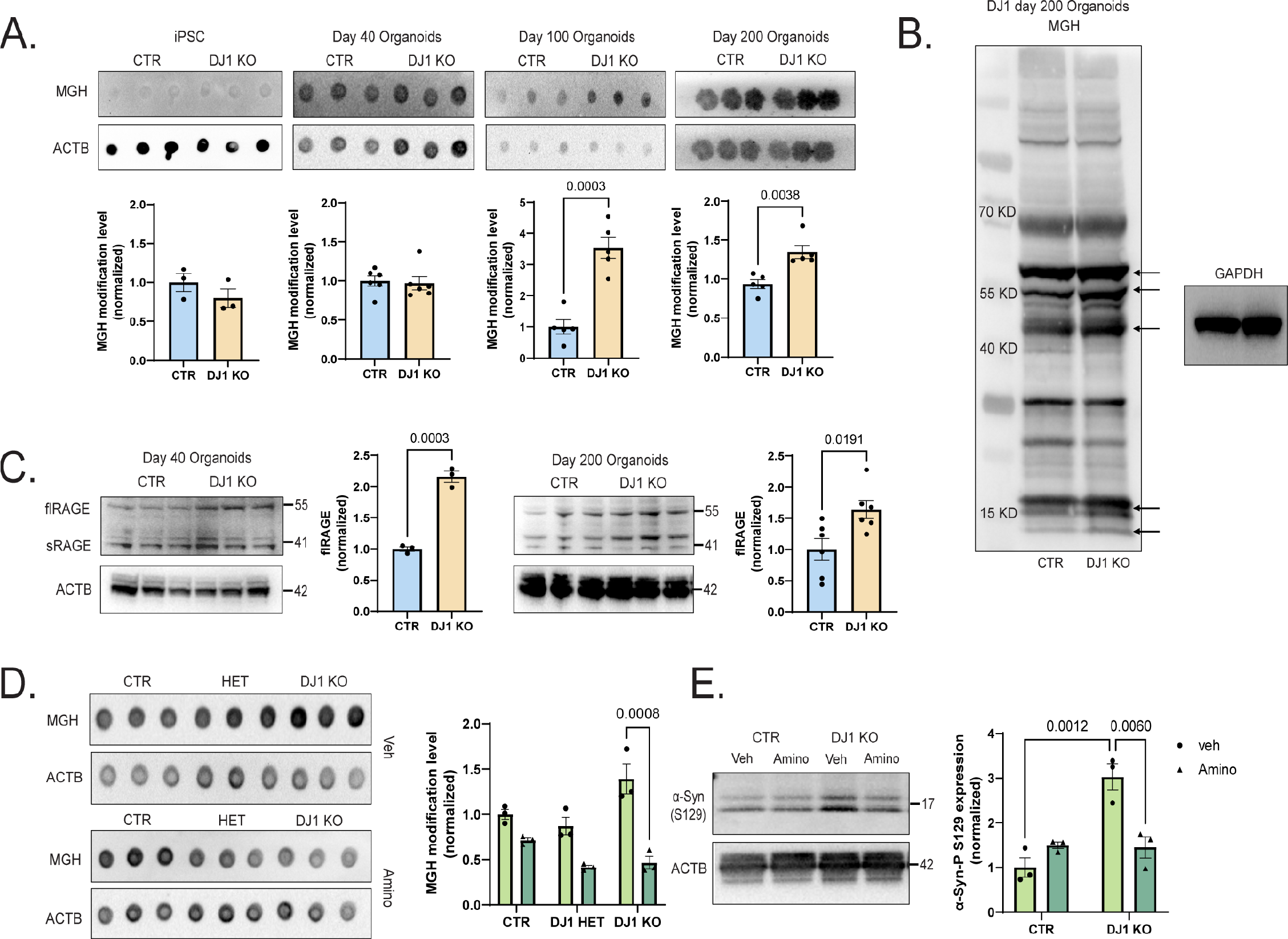
A. Dot blots for MGH protein modification and actin (ACTB)/Ponceau loading control for CTR and DJ1 KO iPSCs, day 40, 100, and 200 midbrain organoids. B. Immunoblot for MGH protein modification CTR and DJ1 KO of day 200 midbrain organoids. C. Immunoblots for RAGE and actin (ACTB) loading control for CTR and DJ1 KO day 40 and day 200 midbrain organoids. D. Dot blots for MGH protein modification and actin (ACTB) loading control in vehicle (Veh) or aminoguanidine (Amino) treated CTR and DJ1 KO day 200 midbrain organoids. E. Immunoblots fot phospho-α-syn (S129) and actin (ACTB) loading control in vehicle (Veh) or aminoguanidine (Amino) treated CTR and DJ1 KO day 200 midbrain organoids. All data are represented in mean ± S.E.M, data points are individual well differentiation, and the p-value was reported on the graph highlighted comparison.

RAGE expression is a sensitive biomarker for the presence of AGEs (Kierdorf and Fritz, 2013), and we were intrigued to observe significantly increased full-length and soluble RAGE protein in both day 40 and day 200 DJ1 KO midbrain organoids (Figure 2C). The increased levels of RAGE in day 40 organoids could indicate that early glycation damage is below levels of detection by MGH blotting or that non-MGH glycation damage is present. When we used the Seahorse XF assay, we found no change in the glycolysis rate (ECAR) or glycolytic capacity between the hMIDO DJ1KO and control midbrain organoids at day 40 or 100 (Figure S2A). Similarly, mass spectrometry analysis did not reveal any differentially expressed metabolites that correlated with increased reactive glycation or increased glycolysis (Figure S2B). Collectively, these data suggest that the increased AGE observed in DJ1KO midbrain organoids is likely a direct effect of DJ1’s deglycase/GLO activity.

Next, we asked whether AGE accumulation in the midbrain contributes to classical PD phenotypes such as α-synuclein aggregation and lysosomal dysfunction. When we treated day 100 midbrain organoids for forty days with aminoguanidine (Amino), a scavenger of reactive carbonyl groups, we detected a significant reduction of MGH glycated proteins in DJ1 KO midbrain organoids relative to vehicle-treated DJ1 KO organoids (Figure 2D). In parallel with reduced MGH, we also observed significantly decreased α-synuclein phosphorylation (S129) in Amino-treated DJ1 KO organoids (Figure 2E). In contrast, there was no significant difference in α-synuclein phosphorylation in Amino-treated control organoids (Figure 2E). The lack of alterations in DJ1 HETs was expected due to the autosomal recessive nature of the PD *PARK7* mutation. Because toxic reactive dicarbonyls that attack guanine residues in DNA can cause double-strand breaks (Richarme et al., 2017), we measured levels of phospho-H2A.X (P-H2A.X) as an indirect readout for DNA damage response. We observed a consistent increase in the P-H2A.X levels in DJ1 KO iPSCs and hMIDO groups with no alterations in DJ1 HET iPSCs compared to CTR, however no alteration in basal H2A.X levels were found among all the groups (Figure S2C). Taken together, these experiments suggest that AGEs may influence abundance and phosphorylation of α-synuclein.

### DJ1 LOF impairs astrocytic metabolic support of midbrain neurons

Astrocytes are the principal glycolytic cell type in the brain and are therefore at a higher risk of acquiring glycation damage than neurons and other brain cell types. As astrocytes are essential for neuronal homeostasis and degradation of neuronal-derived damaged lipids and proteins such as α-syn (Ioannou et al., 2019; Tsunemi et al., 2020), we next investigated astrocyte function in midbrain organoids. To this end, we developed a protocol for isolating astrocytes from mature midbrain organoids (day 100+) and maintaining them in 2D culture. Isolated astrocytes were immunoreactive for canonical astrocyte markers CD44, EAAT2, S100B, and GFAP (with no significant difference among genotypes) (Figure S3A) and functionally responsive to ATP stimulation measured using the calcium sensor GCaMP7 (Figure S3B). In addition, organoid-derived astrocytes were double positive for the midbrain makers NURR1 and FOXA2 and co-stained with CD49f and CD44 expression, confirming their midbrain identity (Figure S3C).

To investigate non-cell-autonomous effects of DJ1 loss of function mediated by astrocytes, we co-cultured astrocytes with midbrain NPCs that by day 50 expressed TH and featured mature neuronal morphology (Figure 3A). Phospho-α-syn (S129) was present in DJ1 LOF astrocytes groups (L166P-1/L166P-1 and CTR/L166P-1), but at lower levels in L166P-1/CTR and CTR/CTR (Figure 3A). When we analyzed total proteolysis capacity using DQ™ Red BSA, we found that the proteolysis capacity of the DJ1 LOF NPCs was significantly increased when co-cultured with CTR astrocytes (L166P-1/CTR) relative to L166P-1/L166P-1 groups (Figure 3B). In addition, in co-cultures of DJ1 LOF astrocytes with CTR NPCs (CTR/ L166P-1) proteolysis levels were disrupted when compared to the CTR/CTR group (Figure 3B), suggesting that proteolytic clearance may be impaired in DJ1 LOF astrocytes. When we seeded day 100 hMIDO (day 100) with mature DJ1 KO or control astrocytes (Figure 3C) and live-imaged the outgrown of TH-TDtomato^+^ axons and COX8A-emerald^+^ astrocytes to confirm the effectivity of the graft prior to protein collection, we found that the addition of CTR astrocytes to KO midbrain organoids significantly reduced the amount of the late glycation product CML (Figure 3D). In contrast, the addition of DJ1 KO astrocytes did not significantly increase CML levels in CTR midbrain but increased CML 1.6-fold of that in the CTR/CTR.

**Figure 3.**
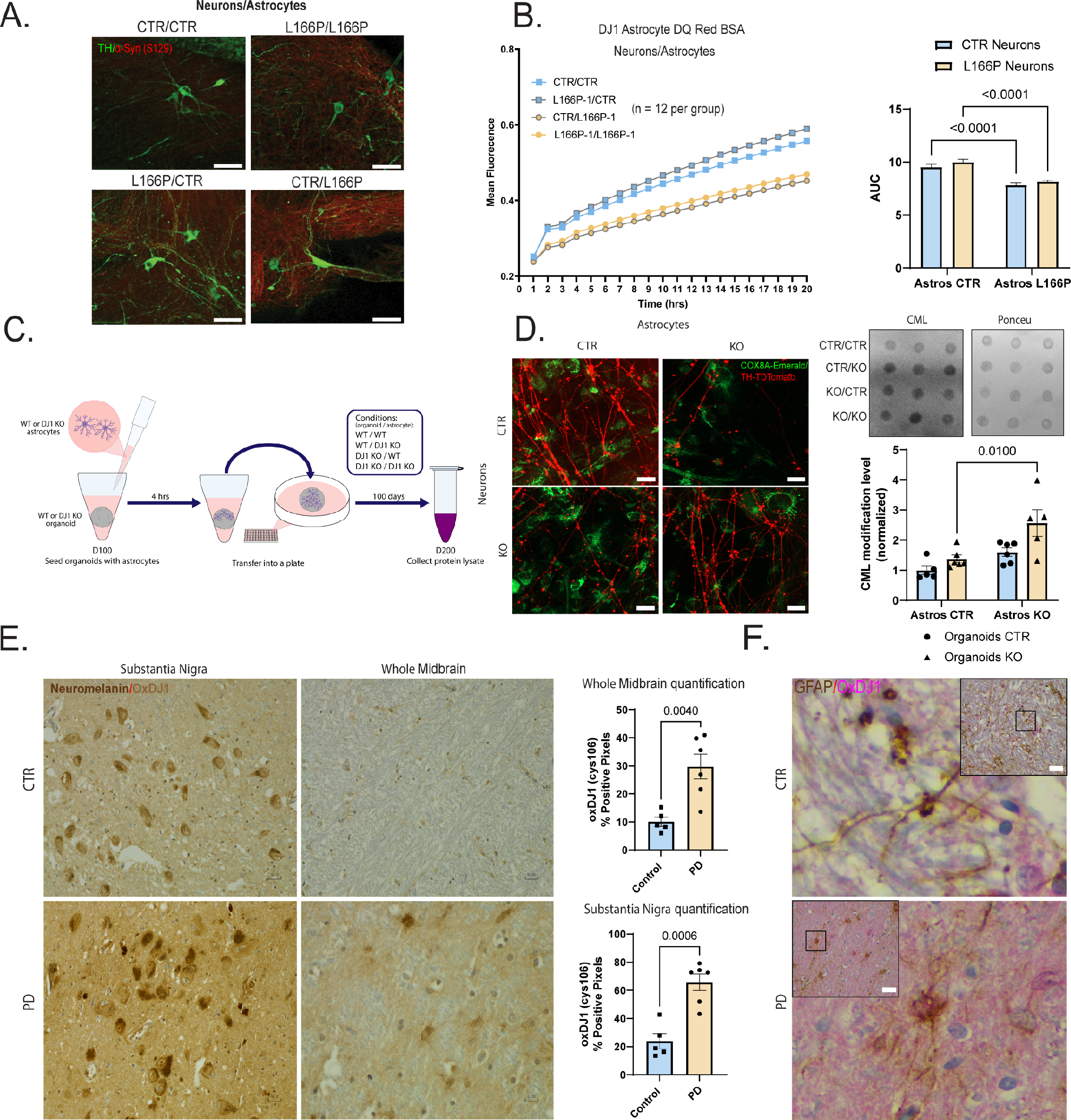
A. 40x images of day 50 NPC/Astrocyte co-cultures stained for TH (green), phospho-α-syn (S129) (Red). B. DQ BSA proteolysis live imaging in mixed genetic neuron/astrocytes co-cultures. C. Astrocytes seeding procedure schematic. D. Astrocytes seeding images of astrocytes following outgrown axons on the left panel. Dot blot for CML protein modification levels in astrocyte seeded organoids and quantification on the right panel. E. 40x micrography of the *substantia nigra* and midbrain of PD patients and age-matched controls showing OxDJ1 staining in light brown. QuPAth of OxDJ1 positive pixel quantification. F. 40x micrography GFAP positive midbrain astrocytes (brown) and OxDJ1 (red). All data are represented in mean ± S.E.M. Scale bars, 15 μm for F.; 50 μm for C., G. and H. data points are individual well differentiation or individual patients, and the p-value was reported on the graph highlighted comparison.

Thus far, our results suggest that loss of DJ1 activity selectively in astrocytes can influence AGE levels in midbrain tissue. To investigate these relationships in the human brain, we quantified cell-type expression of DJ1 in the midbrain of a cohort of PD patients (Table 1). We observed that GFAP-positive astrocytes were positive for ox-cys106 DJ1, which correlates with its increased DJ1 activity. (Figure 3F). We also measured DJ1 activation levels by quantifying ox-cys106 DJ1 in the substantia nigra (SN) and midbrain. When compared to age-matched controls, PD patients had significantly increased levels of ox-cys106 DJ1 in the SN and whole midbrain (Figure 3E, Table 1). Collectively, this analysis suggests that DJ1 has a prominent neuroprotective role in astrocytes and that PD-associated DJ1 LOF variants contribute to neurodegeneration via astrocytes.

### DJ1 LOF astrocytes have a pro-inflammatory phenotype and accumulate aggregated proteins

To identify pathways that contribute to non-cell-autonomous phenotypes mediated by DJ1 LOF astrocytes, we performed global TMT-proteomics and phospho-proteomics in astrocytes derived from 2 clones (L166P-1 and L166P-2) of KOLF 2.1J DJ1 L166P and their respective CTRs. In total, the MS proteomic analysis identified ~8000 different proteins expressed in all samples. PCA analysis showed that the 2 DJ1 L166P clones clustered together and separated from the CTR, and we therefore combined the two DJ1 L166P clones for the final analyses (Figure S4A-B). In total, we identified 1043 differentially expressed proteins in the combined DJ1 L166P dataset (Figure 4A, P-value of <0.01) indicating a significant proteome alteration that included upregulation of inflammatory/reactivity-related proteins such as ANXA3, FGB, AMIGO2, and SERPINE1 and pro-inflammatory interleukins IL32 and IL18 as the most highly differentially upregulated proteins in the DJ1 L166P astrocytes (Figure 4A). When we probed our dataset for PD risk proteins, we found that α-synuclein and UCHL1 were increased, together with the mitochondrial redox balancing enzyme TXNRD2 (Figure 4A).

**Figure 4.**
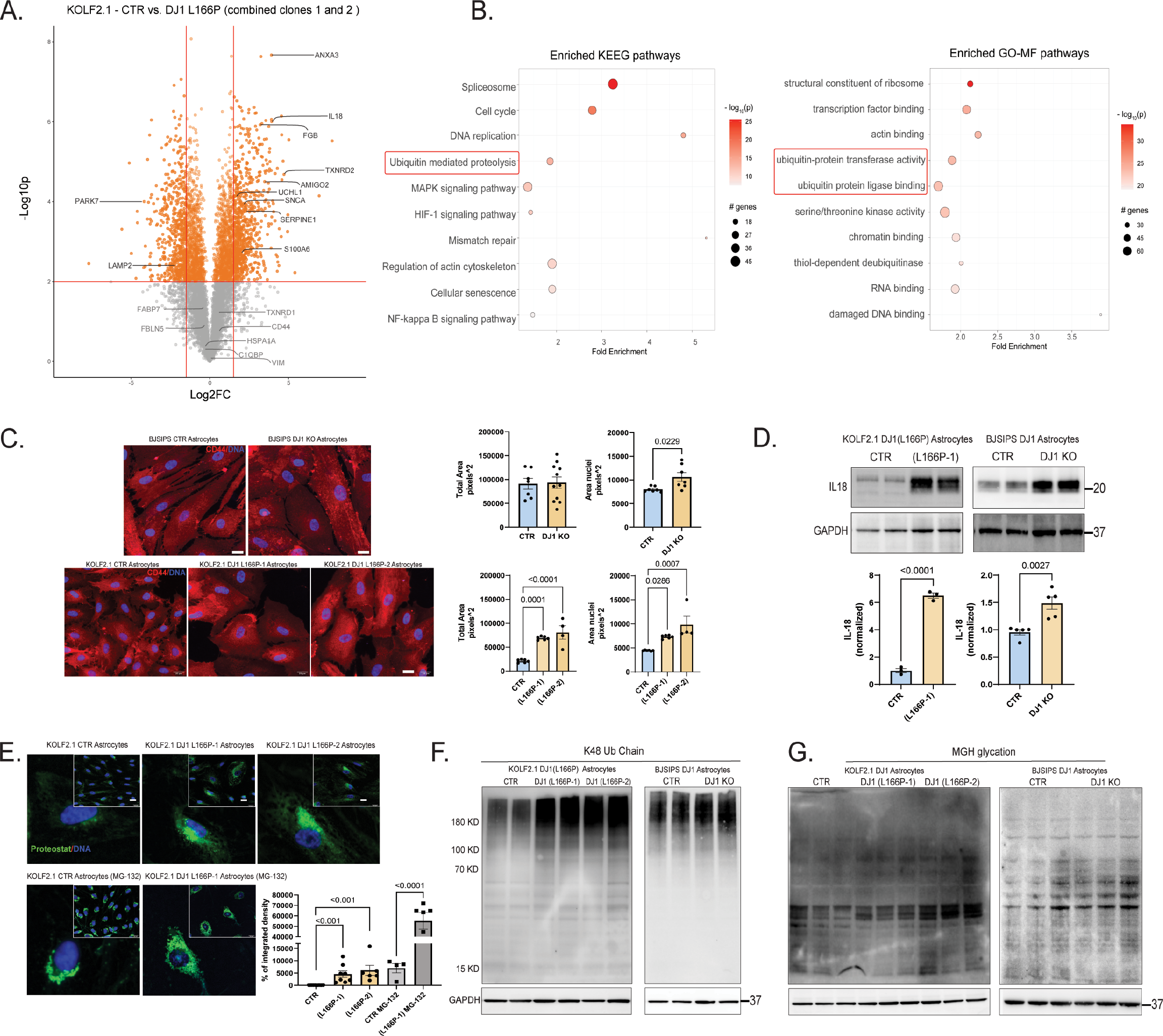
A. Volcano plot representation of TMT-labelled proteomics in KOLF 2.1J and DJ1 L166P KO1 and KO2 midbrain astrocytes with selected proteins labeled. B. KEEG and FO-MF pathways enrichment analysis using pathfinR showing the selected top 10 terms. C. 40x micrography of CD44 (red) and DAPI (blue) staining in BJ-SiPS CTR and KO astrocytes and KOLF 2.1J CTR, DJ1 L166P KO1, and KO2 astrocytes. Quantification of cell total area based on CD44 staining and nuclear area based on DAPI staining. D. IL18 in BJ-SIPS CTR and KO astrocytes and KOLF 2.1J CTR, DJ1 L166P KO astrocytes. E. 40x micrography and quantification proteostat fluorescence levels in KOLF 2.1J CTR, DJ1 L166P KO1 and KO2. F. Immunoblots for K48 ubiquitin chain and G. MGH protein modification. All data are represented in mean ± S.E.M, data points are individual well differentiation, and the p-value was reported on the graph highlighted comparison. Scale bars, 20 μm.

To identify molecular pathways altered in DJ1 L166P astrocytes, we performed enrichment analysis using the Pathfinder package, which considers protein-protein interactions. Among the top terms, we found the ubiquitin/proteasome system (UPS), cytoskeleton modification and inflammatory responses (Figures 4B). We then performed phospho-proteomics to identify differentially expressed phosphorylation sites and predict the most active kinases in DJ1 L166P astrocytes (Figure S4D). Based on this analysis, we identified the CDK isoforms and MAKP8 as the most likely phosphorylating proteins involved in regulating the cytoskeleton and inflammation, including the innate immune response regulator IRAK4 and other related proteins. In addition, when we plotted the CDK family with respect to differentially regulated phosphorylation sites, we observed auto-phosphorylation of CDK1 and increased phosphorylation in cytoskeletal proteins such as MAP4, MAP1B, MAPT, and septin9 (Figure S4D).

Given that reactive astrocytes undergo key morphological change known as hypertrophy (Escartin et al., 2021), we next performed immunostaining for the glial cell body marker CD44 and Hoechst 33342 to evaluate nuclear morphology (Figure 4C). In the full DJ1 KO, we didn’t detect any alterations in the cell body area among the groups, however, the nuclear area was increased in DJ1 KO astrocytes, and both nuclear size and cell body area were increased in DJ1 L166P astrocytes (Figure 4C). We also detected increased IL18 expression in both DJ1KO and DJ1 L166P astrocytes, consistent with our proteomics analysis (Figure 4D).

Our proteomics analysis of DJ1 L166P astrocytes identified alterations in proteolysis and an increase in α-synuclein. To relate these phenotypes with PD pathology, we performed multiple functional assays. In the presence of the proteostat dye (which binds and fluoresces in the presence of protein aggregates), fluorescence intensity was significantly increased in DJ1 L166P astrocytes, indicating an accumulation of protein aggregates (Figure 4E). After treatment with the proteasome inhibitor MG-132, aggregation increased in both DJ1 L166P lines, and CTR astrocytes reached levels comparable to those found in UT L166P (Figure 4E). DJ1 KO and DJ1 L166P astrocytes also expressed higher levels of K48 ubiquitin chain proteins at high molecular weight, characteristic of protein aggregates (Figure 4F). These observations point to a higher UPS activity in DJ1 L166P combined with an inability to degrade protein aggregates. When we integrated the proteomics data with the aggregation risk scores (generated ZaggSC and TANGO (Ciryam et al., 2013)) for each differential protein, we identified several proteins enriched in DJ1 L166P cells that were at increased risk for neurodegeneration-associated aggregation such as APOA1, JADE1, and SERP1 (Figure S4C). In addition, similar to DJ1 KO hMIDOs, DJ1 LOF in astrocytes also led to increased levels of MGH-glycated proteins for specific molecular weight bands, indicating that the phenotype was biased to certain vulnerable proteins (Figure 4G). Altogether, these analyses suggest that proteome instability leads to inflammation and reactivity, and consequently to cytokine release in astrocytes.

### DJ1 loss of function impairs the lysosome, leading to accumulation of α-syn monomers and oligomers in astrocytes

Degradation of aggregated proteins, defined as autophagic flux (the rate of autophagic degradation) is upregulated in early stages of neurodegenerative diseases (Bordi et al., 2016; Fleming et al., 2022). Next, we sought to determine whether the accumulation of protein aggregates observed in DJ1 LOF astrocytes resulted from impaired autophagy or reduced ability to increase autophagy flux. When we analyzed autophagy flux in both astrocytes’ DJ1 LOF lines, we found that the levels of lipidated LC3 II increased significantly in all BAF-treated groups relative to untreated controls (Figure 5A). However, when we compared BAF-treated groups with their respective BAF+ controls, we did not find any alteration in LC3 II levels, (Figure 5A) suggesting that the formation of the autophagosome is not impaired in both DJ1 LOF lines. Next, we analyzed P62 flux to evaluate the ability of the autophagosome to degrade cargo and found that levels of P62 increased significantly for all BAF-treated groups relative to untreated groups (Figure 5A). In contrast with LC3 II, the levels of P62 were increased in the DJ1 L166P KO group and no alteration in the BJSIPS KO groups was found (Figure 5A).

**Figure 5.**
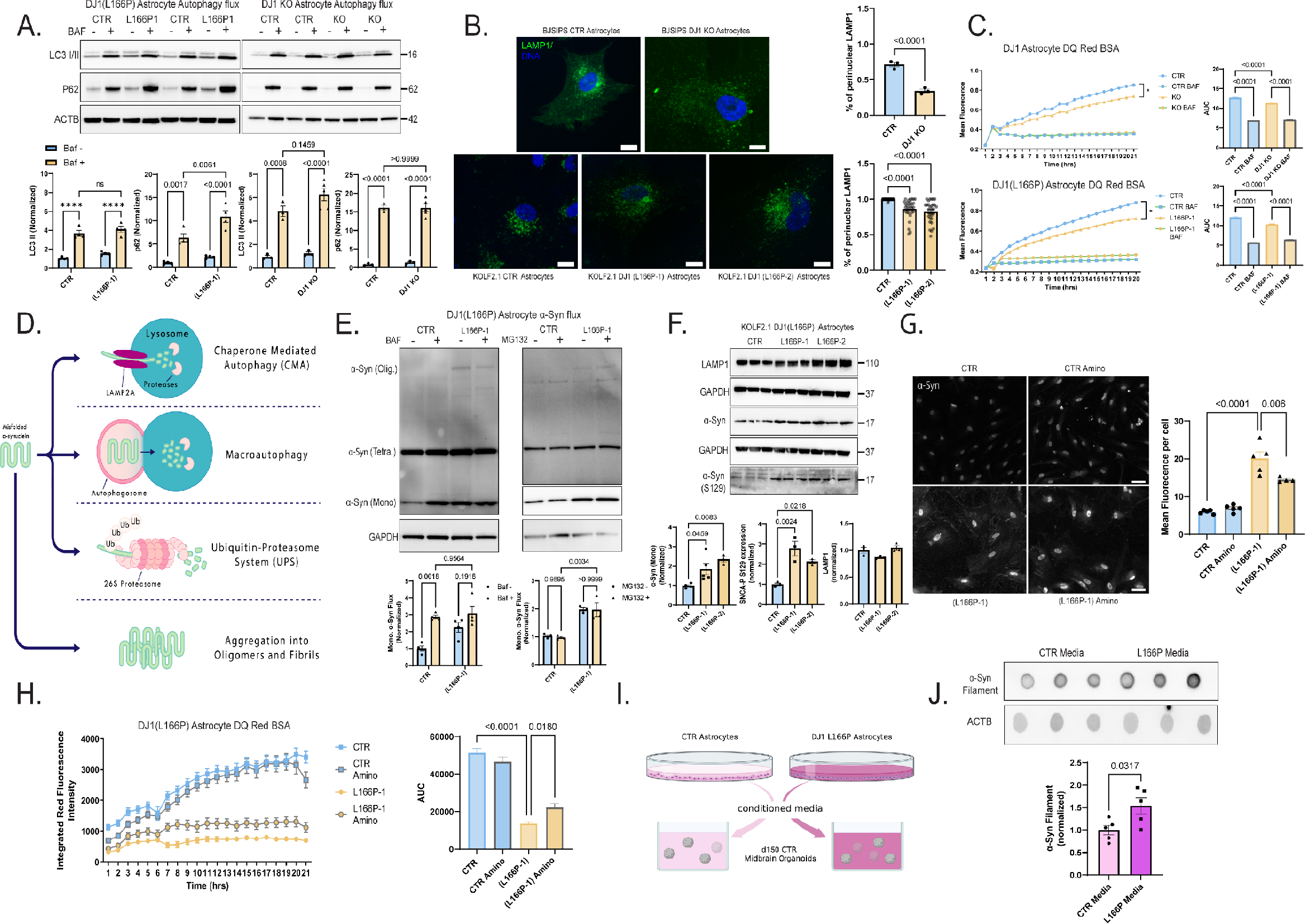
A. Immunoblots for P62, LC3 I/II, actin (ACTB) loading control in BAF – and + treated CTR and DJ1 KO in KOLF 2.1J and BJ-SIPS midbrain astrocytes. B. LAMP1 (green) and DAPI (blue) staining of astrocytes 40x images and quantification of LAMP1 distribution. C. DQBSA proteolysis live imaging assay BAF − and + treated CTR and DJ1 KO KOLF 2.1J astrocytes or CTR and KO BJ-SIPS astrocytes. D. Schematics showing α-syn degradation pathways. E. Immunoblots for α-syn in BAF and MG132 − and + treated CTR and DJ1 KO KOLF 2.1J astrocytes. F. Immunoblots for LAMP1, α-syn, phospho-α-syn (S129), and GAPDH loading control in and KOLF 2.1J CTR, DJ1 L166P KO1, and KO2 astrocytes. G. 40x images of total α-syn (gray) staining of astrocytes and quantification of vehicle (Veh) or aminoguanidine (Amino) treated astrocytes of CTR or DJ1 L166P genotypes. H. DQBSA proteolysis live imaging assay of vehicle (Veh) or aminoguanidine (Amino) treated astrocytes of CTR or DJ1 L166P genotypes. I. Experimental design for conditional media experiment. J. Dot blots for oligomeric α-syn and actin (ACTB) loading control for CTR and L166P astrocyte media treated midbrain organoids. All data are represented in mean ± S.E.M, data points are individual well differentiation, and the p-value was reported on the graph highlighted comparison. Scale bars, for B-15 μm; H-40 μm.

Next, we assessed perinuclear localization of the lysosome, an essential mechanism for chaperone-mediated autophagy (CMA) activity and protein degradation (Kiffin et al., 2004). In both DJ1 KO and DJ1 L166P astrocytes, we observed a decrease in perinuclear endo/lysosomes, as detected by LAMP1 and DAPI immunostaining, relative to the CTR group (Figure 5B). Consistently, when we quantified proteolytic function using live imaging of DQ™ Red BSA, which releases fluorescence upon proteolysis, we detected impaired proteolysis in both DJ1 KO and L166P astrocytes (Figure 5C). Inhibition of the lysosome with BAF decreased fluorescence intensity, demonstrating that BSA proteolysis is largely performed by the lysosome (Figure 5C). To deepen our understanding of the impaired lysosomal function, we next evaluated presence of the early and late lysosomal damage markers GAL3 and K48, and the repair marker CHM4b (Radulovic et al., 2018). In the DJ1 L166P KO groups, percentages of LAMP1-positive puncta that were also positive for GAL3 and K48 were both increased, and we also observed an increase in K48 intensity per area relative to control (Figure S5A). The percentage of LAMP1-positive puncta for the repair marker CHMP4b also increased in the DJ1 L166P KO groups (Figure S5A). Collectively, these data reveal that although the endo/lysosomal system in DJ1 L166P KO cells sustains more damage, it is still capable of triggering repair systems.

Toxic soluble and insoluble forms of α-syn are believed to accumulate in midbrain cells when degradation of aggregated forms through macro-autophagy fails to meet the demand of the elevated α-syn flux. α-Syn degradation is primarily performed by CMA and with the UPS as the preferred mechanism for aggregated forms. (Figure 5D). We analyzed the autophagy capacity to degrade monomeric α-syn, and found that monomeric α-syn in BAF-treated CTR increased whereas the DJ1 L166P KO group had a decreased flux (Fig 5E). Untreated DJ1 KO astrocytes also had significantly higher synuclein monomers compared to control astrocytes (Figure 5E). In contrast, the amount of synuclein monomers in BAF-treated control astrocytes did not decrease, suggesting that DJ1KO impairs the lysosomal clearance of synuclein monomers (Figure 5E). In addition, α-syn oligomers were the main form present in DJ1 L166P KO (Figure 5E). Although the UPS is known to contribute to α-syn degradation (Song et al., 2009a), levels of α-syn did not significantly increase when control or DJ1KO astrocytes were treated with the proteasomal inhibitor MG132, suggesting that proteasomal activity has minimal contribution to a-syn degradation in astrocytes (Figure 5E). In addition, the monomeric unmodified and S129 phosphorylated forms of α-syn increased in the DJ1 L166P lines compared to the CTR, while no alteration was observed in the endo/lysosome marker LAMP1 (Figure 5F).

Based on our data thus far, we hypothesized that AGE damage is at least partly responsible for the reduced lysosomal proteolysis observed in DJ1 LOF astrocytes. When we treated astrocytes with the Methylglyoxal (MGO) scavenger Amino for 10 days, we noticed that lysosome proteolysis was significantly improved in L166P KO astrocytes (Figure 5H). Consistently, total α-syn levels were significantly reduced in treated relative to untreated KO L166P cells with no differences observed in treated controls (Figure 5G). These data indicate that Amino specifically relieves glycation damage in KO L166P cells. To evaluate glycation stress more specifically, we treated DJ1 KO astrocytes with the reactive dicarbonyl MGO, the main cause of glycation in live cells, and measured apoptotic nuclei and poly-caspase activation. Relative to untreated control cells, we detected significantly higher numbers of apoptotic nuclei and higher levels of poly-caspase in untreated DJ1 KO astrocytes (Figure S3D). In addition, MGO treatment significantly increased the number of apoptotic nuclei and active caspase in DJ1 KO, while no change in cell death hallmarks was observed in CTR astrocytes (Figure S3D). When we examined proteolysis, we found decreased levels in DJ1 L166P astrocytes, and that these levels were even further reduced by MGO treatment (Figure S3E). To identify weather soluble factors release by the DJ1 LOF astrocytes could be causing the toxic effect on neurons, we performed a conditioned media experiment (Figure 5I). We detected significant increase in soluble α-syn oligomers in hMIDOs treated with L166P astrocyte media, which indicates release of toxic soluble factors by L166P astrocytes (Figure 5J). Taken together, our data show that DJ1 loss of function reduces lysosomal capacity, resulting in the accumulation of toxic forms of synuclein and increased neuronal cell death.

## DISCUSSION

Failed protein quality control and proteome damage are common features of several neurodegenerative diseases (Ahfeldt et al., 2020; Bourdenx et al., 2021; Burbulla et al., 2017). Here, we show that DJ1 activity is essential to prevent accumulation of toxic damaged proteins in the midbrain. Although we did not detect alterations in soluble α-syn monomers as a consequence of DJ1 LOF, we detected an increase in soluble oligomers, indicative of dynamic relationships between α-syn oligomers and oligomerized or aggregated proteins that may be routed via different degradation processes. At a cellular level, our data show that DJ1 LOF or mutations hinder lysosomal processing of α-syn in astrocytes. Consequently, soluble α-syn oligomers aggregate into toxic forms, resulting in direct oxidative damage to the proteome of midbrain astrocytes and subsequent activation of astrocyte-derived inflammation signals that further aggravate pathology. These data deepen and extend from the established view of astrocytes in PD inflammation.

Glycation is a form of oxidative protein damage commonly observed in diabetes, a risk factor for developing PD (Konig et al., 2018; Yang et al., 2017). We observed accumulation of MGH, an early form of glycation damage derived from MGO, in non-aggregated proteins in both hMIDOs and isolated astrocytes. Nonetheless, non-aggregated glycated forms are thought to be degraded by the proteasome, and build-up can trigger ER stress (Adamopoulos et al., 2014; Raupbach et al., 2020). In what way that DJ1 protects from protein glycation damage is still unclear. Early work proposed that DJ1 integrates with the glycation stress system through enzymatic function (Hasim et al., 2014), participates in MGO degradation, or prevents permanent glycation damage by direct repair of early glycation products (Richarme et al., 2015, 2017). In patient derived iPSC neurons bearing DJ1 mutation glycation related products were found elevated (Mazza et al., 2022). However, DJ1 deficiency fail to enhance neuronal cell death when challenged with MGO (Mazza et al., 2022). In the DJ1 KO organoids, we observe a progressive accumulation of glycated proteins starting at day 100. Glycolysis levels and MGO precursor metabolites were both unaltered, which altogether points to a failure in MGO degradation or protein glycation repair. Conversely, presence of the carbonyl scavenger Amino decreased levels of MGH-modified proteins in both CTRs and DJ1 KO, validating its ability to prevent glycation stress, potentially by preventing the glycation stress-induced increase in α-syn phosphorylation. Altogether, these findings suggest that DJ1 participates in glycation metabolism and DJ1 mutation contribute to PD by causing slow accumulation deleterious glycation products.

In the absence of DJ1, levels of inflammatory cytokines IL18 and IL32 and downstream signal events such as IRAK4 phosphorylation both increased. Consistently, the proteome analysis identified various upregulated proteins such as ANXA3, S100A6, FGB, and SERPINE while others such as VIM and S100A10 were unchanged in DJ1 LOF astrocytes. These data reveal a unique astrocyte signature with relevance for PD-associated pathology, consistent with recent findings that reported glial cell activation upon scRNA-seq analysis of the human midbrain in PD patients (Smajić et al., 2022). In a broad number of cells including astrocytes, cytoskeleton modification and changes in morphological features to sense, mobilize, or invade are the main characteristics of inflammation. The proteome and phosphorylation pattern also indicates an elevated modification of microtubule-associated proteins, which can explain the altered morphology of the affected astrocytes and the increase in nuclear size and cell body hypertrophy we observed in cells with the L166P mutation. Consistently, we found differential expression of MAP4, MAP1B, MAPT, and other proteins related to cytoskeleton modification. Additionally, our prediction analysis identified the CDK kinase family, reported to phosphorylate cytoskeleton-associated proteins and other related proteins (Sánchez et al., 2000), as the top active kinases predicted to phosphorylate the enriched residues.

Many PD genes participate in convergent pathways that are altered in the pathology (Kumaran and Cookson, 2015). Therefore, the observation that additional familial PD genes, such as *SNCA* and *UCHL1*, were upregulated in LOF DJ1 astrocytes corroborated these observations. We observed this phenotype in isolated astrocytes extracted from organoids, which points to the crucial contribution of astrocytes to specific PD phenotypes, such as α-syn accumulation. However, we cannot exclude a previously reported prion-like acquisition in which the spreading of α-syn throughout the hMIDOs causes damage (Bernis et al., 2015).

In addition to lysosomal changes, we also identified UPS system alterations in our enrichment pathway analysis, consistent with the increase in aggregated protein inclusions and the sensitivity of the DJ1 mutation cell lines to proteasome inhibition. These relationships indicate the presence of ongoing protein damage that continuously needs to be repaired by the proteasome and lysosome systems. Alterations in autophagy were relatively mild in our DJ1 L166P LOF astrocytes but more pronounced in the DJ1 KO. Although we only detected a mild alteration of autophagy in the L166P lines, α-syn oligomers were formed as a consequence of impaired lysosomal degradation of α-syn. These relationships indicate a varied phenotype depending on the type of mutation, with the DJ1 L166P mutation generating the most deleterious phenotypes as it generates an inactive DJ1 protein. Truncated DJ1 misfolds and therefore constantly needs to be degraded by the UPS system which could further exacerbate the disease-associated phenotypes (Miller et al., 2003; Olzmann et al., 2004). In agreement, we found an increase in ATF6, which has been shown to have increased activation due to the accumulation of misfolded products, leading to the induction of inflammatory gene networks (Rao et al., 2014).

Dysregulated networks common to early-onset and sporadic PD converge on proteostasis failure, causing accumulation of aggregated proteins (Ahfeldt et al., 2020; Burbulla et al., 2017). Based on our studies in the early-onset PD DJ1 model, we hypothesize that accumulation of glycated products, which overburden and damage proteins degradation and repair systems, ultimately leads to proteostasis failure. In addition, due to their high non-aerobic glycolysis, astrocytes would be particularly vulnerable to this oxidative damage. In conclusion, we provide evidence that glycation and widespread protein aggregation are fundamental phenotypes in familial DJ1-linked PD, pointing to strategies for developing more effective therapeutics.

## STAR+METHODS

### Resource availability

#### Lead contact

Further information and requests for resources and reagents should be directed to and will be fulfilled by the lead contacts, Joel Blanchard and Tim Ahfeldt.

### Materials availability

All resources and materials reported in this paper will be shared by the lead contact upon request.

### Data and code availability

The MS proteomics data have been deposited to the MassIVE repository with the dataset identifier MSV000090202.

Reviewer access: ftp://MSV000090202@massive.ucsd.edu Password: DJ1_Parfitt

The metabolomics is submitted at the NIH Common Fund’s National Metabolomics Data Repository (NMDR) website, the Metabolomics Workbench, https://www.metabolomicsworkbench.org where it has been assigned Project ID (not assigned). The data can be accessed directly via it’s Project DOI: (not assigned).

### Generation of isogenic knockout lines

Human iPSCs were cultured in standard conditions, and before nucleofection cells were pre-treated for 1 h. with 10 μM Rhok inhibitor. 4×10^6^ cells were dissociated using Accutase. Cells were pelleted and resuspended in 800 μl PBS^−/−^ containing 5 μg px330 CRISPR DNA each and transferred into nucleofection cuvettes. Nucleofection was carried out using either the P3 Nucleofector kit from Amaxa and the standard and program CB-150 or the primary P4 Nucleofector kit from Amaxa and the standard and program hiPSC CA-137. The iPSC lines used for the generation of the DJ1 KO were from the BJSIPS background and the guides as described in (Ahfeldt et al., 2020). The iPSC lines used for the generation of the DJ1 L166P point mutation were from KOLF 2.1J background (Pantazis et al., 2021). The parental cell lines were karyotyped prior to the begging of the experiments. Genotyping PCR was used to identify clones with homozygous or compound heterozygous deletions leading to truncations and frameshift mutations. Clones for all lines containing deletions were identified by Sanger sequencing.

### Midbrain differentiation and organoid maintaining

hiPSCs were cultured in Stemflex™ medium (ThermoFisher) at 37°C, with 5% CO2 in a humidified incubator, as previously described (Sarrafha et al., 2021). For the organoid aggregation and differentiation, 125-ml disposable spinner flasks (Corning, VWR) were placed on a nine-position stir plate (Dura-Mag) at a speed of 65 rpm, as previously reported, starting with dissociated 40×10^6^ hPSCs in Stemflex + Rhok inhibitor Y-27632 (4 μM). Differentiation was initiated when spheres reached 300-500 μm by dual-SMAD inhibition with SB431542 (R&D Systems, 10 μM), LDN193189 (Stemgent, 100 nM), B27-Vit A and N2 in DMEM-F12, to unsure the proper size range spheres were filtered using a set of 300 and 500 μm filters (pluriSelect). Midbrain-specific patterning for midbrain NPCs organoids was the addition of CHIR99021 (Stemgent, 3 μM), Purmorphamine (STEMCELL, 2 μM), and SAG (Abcam, 1 μM) (Kriks et al., 2011). Post patterning Neural maturation medium was DMEM F12 medium containing N2, B27-VitA, 20 ng/mL GDNF (R&D Systems), 20 ng/mL BDNF (R&D Systems), 0.2 mM ascorbic acid (Sigma), 0.1 mM dibutyryl cAMP (Biolong), 10 μM DAPT (Cayman Chemical). For long-term maintenance (after day 35), the spheres were transferred to ultralow attachment plates (Corning, VWR) at 5 spheres per ml of media. The medium for long-term culture was DMEM F12 medium containing N2, B27-VitA, 10 ng/ML GDNF (R&D Systems), 10 ng/mL BDNF (R&D Systems), 0.2 mM ascorbic acid (Sigma).

### Astrocyte differentiation and isolation

We derived astrocytic 2D cultures from large-scale 3D spin cultures, starting at day 90 using a protocol adapted from (TCW et al., 2017). This involved dissociation and serial passaging of midbrain organoids under conditions that favor astrocyte growth. Cells are grown in 125 ml flasks containing hundreds of individual organoids totaling more than 4×10^8^ cells. At this time, the culture is comprised of various midbrain cell types including DNs, other neurons, progenitor cells, and astrocytic cell types. To isolate astrocytic progenitors, we gently triturated organoids in trypsin enzyme solution using a glass pipette to break up individual organoids into large chunks. After washing steps and pelleting of the organoids, we plated the suspension on 15 cm dishes coated with 0.1% gelatin. Cells are maintained in an astrocyte propagation medium (Astrocyte Medium, ScienCell #1801) for a week or until the first astrocytes are attached and start to divide. Cells were passaged to a maximum of P3. The maintaining and experimental media consisted of Advanced DMEM/F-12 (1 part, ThermoFisher 12634010), Neurobasal (1 part, ThermoFisher 21103049), B-27 Supplement, N-2 Supplement, MEM Non-Essential Amino Acids Solution (ThermoFisher 11140050), GlutaMAX (ThermoFisher 35050061), CNTF (10 ng/ml, PeproTech 450-13). For passaging, Astrocyte Medium was used on the first day and replaced by Astrocyte mature after 70% of confluency was reached. For the conditioning media experiments, mature media added to a confluent plate of astrocytes. After 3 days the media was collected, filtered and then added to day 150 hMIDOs.

### Midbrain organoid astrocyte seeding and co-culture

For the co-culture experiments midbrain NPCs of day 14 after differentiation and day 100 astrocytes were plated in a 96 well plate in a 5:1 proportion. They were maintained in a post patterning neural maturation medium for 2 weeks and then transferred to a long-term culture medium for the experimental procedures. For the seeding procedure, 40 organoids were seeded with 50k astrocytes in a V bottom plate (Corning). After a week, organoids were transferred to a 96 well plate 1 organoid per well and maintained until day 200. The efficiency of the seeding was observed by a GFP reporter present in the exogenous astrocytes.

### Immunocytochemistry of fixed cells

Cells were fixed with 4% PFA for 20 min. Cells were blocked in 0.1% Triton X-100 (Sigma) in 5% horse serum/PBS, and then incubated in primary antibody (table 2) (0.1% Triton X-100 in 5% horse serum/PBS) overnight at 4°C. On the following day, cells were washed in PBS and incubated in species-specific fluorophore-conjugated Alexa fluor secondary antibodies and DAPI nuclear stain according to the manufacturer’s protocol. The imaging was performed using the High content imager CX7 (Thermo Fisher) and phenotyping and quantifications were performed using ImageJ.

### Immunostaining and image analysis of sectioned spheres

Organoids were fixed with 4% PFA O/N and embedded in paraffin. Serial sections (4-6 μm) of paraffin-preserved midbrain organoid sections were prepared using a Leica RM2255 microtome, sections were placed on charged slides and baked overnight at 70°C. IHC was performed on Ventana Benchmark XT. Antigen retrieval with CC1 (citric acid buffer) was performed for 1 h, followed by primary antibody incubation for 30 minutes (min.). A multimer secondary antibody was used for all samples. IHC sections were imaged using an Aperio VERSA 8 (Leica Biosystems, Wetzlar Germany) digital slide scanner and analyzed in QuPath (version 0.2.3, https://QuPath.github.io/).

### Immunohistochemistry of human tissue

Cases and controls brain samples were derived from the Mount Sinai Neuropathology Brain Bank. Inclusion criteria were individuals with a neuropathological diagnosis of Parkinson’s disease for cases and cognitively normal with no or only age-related neuropathological changes for controls. Formalin-fixed paraffin-embedded (FFPE) sections (5μm) were prepared from substantia nigra blocks, mounted on positively charged slides, and baked overnight at 70° C. Immunohistochemistry (IHC) was performed on a Ventana Benchmark XT automatic staining platform (Roche Diagnostics, Indianapolis, IN) according to the manufacturer’s protocol with reagents and antibodies acquired from the same lot. Antigen retrieval was done using CC1 buffer (Tris/Borate/EDTA buffer, pH 8.0-8.5, Roche Diagnostics) for 1 h followed by primary antibody incubation. All primary antibodies were diluted in antibody dilution buffer (ABD24, Roche Diagnostics). Primary antibodies were incubated for 36 min (mOXDJ1, 1:400, Abcam) or 28 min (GFAP, 1:10, Ventana, 760-4345) followed by either 3,3’-diaminobenzidine (DAB) or alkaline phosphatase for visualization. For slides that were double-labeled, both DAB and alkaline phosphatase were used for visualization. All slides were counterstained with hematoxylin and coverslipped.

### Digital Histopathologic Analysis

For unbiased digital quantitative assessment, slides were imaged using an Aperio VERSA 8 (Leica Biosystems, Wetzlar Germany) digital slide scanner. Whole tissue sections and the substantia nigra were manually neuroanatomically segmented on whole slide images (WSI) and analyzed in QuPath (version 0.2.3, https://QuPath.github.io/). All analysis was batch-processed using a custom positive pixel-based analysis workflow that measured the percentage of positive pixels detected using a positive pixel classifier based on thresholded values for DAB intensity. All quantitative values were normalized to the area.

### Live-cell imaging to access lysosome function

Astrocyte monocultures or midbrain neuronal/astrocyte co-cultures were plated in the 96 well plates. Cells were incubated with DQ™ Red BSA, which emits red fluorescence upon proteolysis, for 30 min, washed 3 times with PBS, and kept in maintaining media throughout the assay. Bafilomycin A2 (100 nM, Sigma-Aldrich B1793) was added to specific wells to confirm the lysosomal nature of the proteolysis. Aminoguanidine (30 μM) was used for the reversal experiments. The plates were incubated for image acquisition in Incucyte® Live-Cell Analysis System for 21h and images were analyzed in the Incucyte® software.

### Calcium imaging in astrocytes

For GCaMP8s imaging, plated astrocytes in 2 cm gelatin-coated plates were placed on a Nikon Eclipse TE2000-U microscope, with a 10X objective. GCaMP8s was excited using a 480 nm (Mic-LED-480A, Prizmatix), an HQ480/40x excitation filter, a Q505LP dichroic mirror, and an HQ535/50m emission filter (Semrock). Fluorescence was projected onto an sCMOS Zyla chamber camera (VSC-01910, Andor), and sampled at a rate of 4.7 fps with a frame exposure of 200 ms at 160×120 pixels (4×4 binning). The light source and sCMOS camera were controlled with the Nikon Elements software (NIS-Elements AR 5.20.01). The astrocytes were continuously perfused during fluorescence recording with ACSF with the following composition (in mM): NaCl 125, KCl 5, D-Glucose 10, HEPES-Na 10, CaCl_2_ 3.1, MgCl_2_ 1.3. The ATP treatment (100 μM) was controlled by a ValveBank8 II (AutoMate Scientific Inc.). ROI segmentation of GCaMP8s, raw fluorescence extraction, and background correction was performed with Nikon Elements software. ΔF/F was calculated using R-studio (R version 4.0.3)

### Western blotting and dot blot

Cell culture lysates were generated using RIPA buffer (Thermo Scientific), 1 or 2% SDS lysis buffer (10 mM tris, 150 mM NaCl, 1mM EDTA) containing protease inhibitor cocktail (Thermo Scientific), and phosphatase inhibitor cocktail (Thermo Scientific). Protein concentration was estimated using the BCA assay (Pierce). For Western blot analysis 20-40 μg total protein was denatured under reducing conditions in 4x Laemmli Sample Buffer (Bio-Rad) by heating for 10 min at 70°C before loading onto a 10% Criterion TGX Precast gel (Bio-Rad), then transferred to a PVDF membrane (0.22 μm; Bio-Rad) using the iBlot 2 dry blotting system (Invitrogen). Membranes were blocked for 1 h at RT in 5% w/v non-fat milk (Santa Cruz) in TBS containing 0.1% v/v Tween-20 (Fisher Scientific; TBS-T). Membranes were then incubated in the indicated primary antibody (in 5% milk/TBS-T) overnight at 4°C, washed 4 times in TBS-T, incubated in species-specific HRP-conjugated secondary antibody (in 5% milk/TBS-T) for 1 h at RT, and then washed 4 times in TBS-T. Membranes were subsequently developed with ECL Western blotting substrate (Pierce). Membranes were then washed once in TBS-T and stripped in stripping buffer (25mM Glycine HCl, pH 2.0, and 1% w/v SDS) with vigorous shaking to remove primary and secondary antibodies, washed three times in TBS-T, and blocked for 1 h (in 5% milk/TBS-T) at RT before probing with the next primary antibody. Dot blots were performed in the same way as western blots but without the gel separation step. The primary antibodies are listed in table 2.

### Proteomics - Cell Lysis and Protein Digestion

At the indicated times, cells were washed twice with ice-cold PBS and snap-frozen. Cell pellets were lysed with in-house RIPA buffer (50 mM HEPES, 150 mM NaCl, 1% sodium deoxycholate, 1% NP-40, 0.1% SDS, 2.5 mM MgCl2, 10 mM sodium glycerophosphate, 10 mM sodium biphosphate) containing in-house protease and phosphatase inhibitor cocktail), to produce whole-cell extracts. Whole-cell extracts were sonicated and clarified by centrifugation (16000×g for 10 min at 4oC) and protein concentrations were determined by the Bradford assay. Protein extracts (40 μg) were subjected to disulfide bond reduction with 5 mM TCEP (room temperature, 10 min) and alkylation with 25 mM chloroacetamide (room temperature, 20 min). Methanol–chloroform precipitation was performed before protease digestion. In brief, four parts of neat methanol were added to each sample and vortexed, one part of chloroform was then added to the sample and vortexed, and finally three parts of water were added to the sample and vortexed. The sample was centrifuged at 8000 rpm for 5 min at room temperature and subsequently washed twice with 100% methanol. Samples were resuspended in 100 mM EPPS pH8.5 containing 0.1% RapiGest and digested at 37°C for 16h with trypsin at a 100:1 protein-to-protease ratio.

### Proteomics - Tandem Mass Tag Labeling

Tandem Mass Tag (TMT and TMTpro) labeling of samples was carried out as followed. For total proteome analysis (40 μg of digested peptides), 8 μL of a 10 μg/μL stock of TMT reagent was added to samples, along with acetonitrile to achieve a final acetonitrile concentration of approximately 30% (v/v). Following incubation at RT for 1 h, the labeling efficiency of a small aliquot was tested for each set, and the reaction was then quenched with hydroxylamine to a final concentration of 0.5% (v/v) for 15 min. The TMT-labeled samples were pooled together at a 1:1 ratio. The total proteome sample was vacuum centrifuged to near dryness and subjected to C18 solid-phase extraction (SPE) (50 mg, Sep-Pak, Waters).

### Proteomics - Off-Line Basic pH Reversed-Phase (BPRP) Fractionation

Dried TMT-labeled sample was resuspended in 100 μl of 10 mM NH4HCO3 pH 8.0 and fractionated using basic pH reverse phase HPLC (Wang et al., 2011). Briefly, samples were offline fractionated over a 90 min run, into 96 fractions by high pH reverse-phase HPLC (Agilent LC1260) through an aeris peptide xb-c18 column (Phenomenex; 250 mm × 3.6 mm), with mobile phase A containing 5% acetonitrile and 10 mM NH4HCO3 in LC-MS grade H2O, and mobile phase B containing 90% acetonitrile and 10 mM NH4HCO3 in LC-MS grade H2O (both pH 8.0). The 96 resulting fractions were then pooled in a non-continuous manner into 24 fractions used for subsequent mass spectrometry analysis. Fractions were vacuum centrifuged to near dryness. Each consolidated fraction was desalted via Stage Tip, dried again via vacuum centrifugation, and reconstituted in 5% acetonitrile, and 1% formic acid for LC-MS/MS processing. For Phospho-peptides, dried peptides were fractionated according to the manufacturer’s instructions using High pH reversed-phase peptide fractionation kit (Thermo Fisher Scientific) for the final 6 fractions and subjected to C18 StageTip desalting prior to MS analysis.

### Proteomics - Fe2+-NTA Phosphopeptide Enrichment

Phosphopeptides were enriched using Pierce High-Select Fe2+-NTA phosphopeptide enrichment kit (Thermo Fisher Scientific, A32992) following the provided protocol. In brief, dried peptides were enriched for phosphopeptides and eluted into a tube containing 25 μL 10% formic acid (FA) to neutralize the pH of the elution buffer and dried down. The unbound peptides (flow-through) and washes were combined and saved for total proteome analysis.

### Proteomics – Total proteomics analysis

Mass spectrometry data were collected using an Orbitrap Eclipse Tribrid mass spectrometer (Thermo Fisher Scientific, San Jose, CA) coupled to an UltiMate 3000 RSLCnano system liquid chromatography (LC) pump (Thermo Fisher Scientific). Peptides were separated on a 100 μm inner diameter microcapillary column packed in-house with ~30 cm of HALO Peptide ES-C18 resin (2.7 μm, 160 Å, Advanced Materials Technology, Wilmington, DE) with a gradient consisting of 5%–23% (0-75 min), 23-40% (75-110min) (ACN, 0.1% FA) over a 120 min run at ~500 nL/min. For analysis, we loaded 3/10 of each fraction onto the column. Each analysis used TMT-MS2 based quantification, combined with the FAIMS Pro Interface (using previously optimized 3 CV parameters for TMT multiplexed samples (Schweppe et al., 2019). The scan sequence began with an MS1 spectrum (Orbitrap analysis; resolution 120,000 at 200 Th; mass range 400−1500 m/z; automatic gain control (AGC) target 4×105; maximum injection time 50 ms).

Precursors for MS2 analysis with desired charge state (z: 2-6) were selected using a cycle type of 1.25 sec/CV method (FAIMS CV=−40/−60/−80). MS2 analysis consisted of high energy collision-induced dissociation (HCD) and was analyzed using the Orbitrap (resolution 50,000 at 200 Th; NCE 38; AGC 2×105; isolation window 0.5 Th; maximum injection time 172 ms). Monoisotopic peak assignment was used, precursor fit filter was used (80% fit) and previously interrogated precursors were excluded using a dynamic window (150 s ±10 ppm). For TMTpro analysis, a similar setup was used with the following modifications. Each analysis used Multi-Notch MS3-based TMT quantification (McAlister et al., 2014), combined with a newly implemented Real Time Search analysis software (Erickson et al., 2019; Schweppe et al., 2020). MS2 analysis consisted of collision-induced dissociation (quadrupole ion trap analysis; Rapid scan rate; AGC 1.0×104; isolation window 0.5 Th; normalized collision energy (NCE) 35; maximum injection time 35 ms). Monoisotopic peak assignment was used, precursor fit filter was used (70% for a fit window of 0.5 Th) and previously interrogated precursors were excluded using a dynamic window (180 s ±10 ppm). Following the acquisition of each MS2 spectrum, a synchronous-precursor-selection (SPS) API-MS3 scan was collected on the top 10 most intense ions b or y-ions matched by the online search algorithm in the associated MS2 spectrum (Erickson et al., 2019; Schweppe et al., 2020). MS3 precursors were fragmented by high energy collision-induced dissociation (HCD) and analyzed using the Orbitrap (NCE 45; AGC 2.5×105; maximum injection time 200 ms, the resolution was 50,000 at 200 Th). The closeout was set at two peptides per protein per fraction so that MS3s were no longer collected for proteins having two peptide-spectrum matches (PSMs) that passed quality filters.

### Proteomics – Phospho-proteomics analysis

Mass spectrometry data were collected using an Orbitrap Eclipse Tribrid mass spectrometer (Thermo Fisher Scientific, San Jose, CA) coupled to an UltiMate 3000 RSLCnano system liquid chromatography (LC) pump (Thermo Fisher Scientific). Peptides were separated on a 100 μm inner diameter microcapillary column packed in-house with ~30 cm of HALO Peptide ES-C18 resin (2.7 μm, 160 Å, Advanced Materials Technology, Wilmington, DE) over a 155 min run at ~500 nL/min. For analysis, we loaded half of each fraction onto the column. Each analysis used the FAIMS Pro Interface (using previously optimized 3 CV parameters for TMT-labeled phosphopeptides) to reduce ion interference. The scan sequence began with an MS1 spectrum (Orbitrap analysis; resolution 120,000 at 200 Th; mass range 350−1500 m/z; automatic gain control (AGC) target 4×105; maximum injection time 50 ms). Precursors for MS2 analysis were selected using a cycle type of 1.25 sec/CV method (FAIMS CV=−40/−60/−80). MS2 analysis consisted of high energy collision-induced dissociation (HCD) (Orbitrap analysis; resolution 50,000 at 200 Th; isolation window 0.5 Th; normalized collision energy (NCE) 38; AGC 2×105; maximum injection time 172 ms). Monoisotopic peak assignment was used, precursor fit filter was used (80% for a fit window of 0.5 Th) and previously interrogated precursors were excluded using a dynamic window (120 s ±10 ppm)(Schweppe et al., 2020).

### Proteomics – Data analysis

Mass spectra were processed using a Comet-based (2019.01 rev. 5) software pipeline (Eng et al., 2013). Spectra were converted to mzXML and monoisotopic peaks were re-assigned using Monocle (Rad et al., 2021). MS/MS spectra were matched with peptide sequences using the Comet algorithm along with a composite sequence database including the Human Reference Proteome (2020-01 - SwissProt entries only) UniProt database, as well as sequences of common contaminants. This database was concatenated with one composed of all protein sequences in the reversed order. Searches were performed using a 50 ppm precursor ion tolerance for analysis. TMT or TMTpro tags on lysine residues and peptide N termini (+229.162932 TMT; (+304.207 Da TMTpro) and carbamidomethylation of cysteine residues (+57.021 Da) were set as static modifications, while oxidation of methionine residues (+15.995 Da) was set as a variable modification. For the phosphorylation dataset search, phosphorylations (+79.966 Da) on Serine or Threonine were set as additional variable modifications. Peptide-spectrum matches (PSMs) were adjusted to a 1% false discovery rate (FDR) (Elias and Gygi, 2007). PSM filtering was performed using a linear discriminant analysis(Huttlin et al., 2010), while considering the following parameters: XCorr (or Comet Log Expect), ΔCn (or Diff Seq. Delta Log Expect), missed cleavages, peptide length, charge state, and precursor mass accuracy. For protein-level comparisons, PSMs were identified, quantified, and collapsed to a 1% peptide false discovery rate (FDR) and then collapsed further to a final protein-level FDR of 1% (Savitski et al., 2015). Moreover, protein assembly was guided by principles of parsimony to produce the smallest set of proteins necessary to account for all observed peptides. For TMT-based reporter ion quantitation, we extracted the summed signal-to-noise (S:N) ratio for each TMT channel and found the closest matching centroid to the expected mass of the TMT reporter ion (integration tolerance of 0.003 Da). Reporter ion intensities were adjusted to correct for the isotopic impurities of the different TMT reagents according to manufacturer specifications. Proteins were quantified by summing reporter ion signal-to-noise measurements across all matching PSMs, yielding a ‘‘summed signal-to-noise’’ measurement. For total proteome, PSMs with poor quality, MS3 spectra with 6 or more TMT reporter ion channels missing, or isolation specificity less than 0.8, or with TMT reporter summed signal-to-noise ratio that was less than 150 or had no MS3 spectra were excluded from quantification. Phosphorylation site localization was determined using the AScore algorithm. AScore is a probability-based approach for high-throughput protein phosphorylation site localization. Specifically, a threshold of 13 corresponded to 95% confidence in site localization.

Protein or peptide quantification values were exported for further analysis in Microsoft Excel, R package, and Perseus. Each reporter ion channel was summed across all quantified proteins and normalized assuming equal protein loading of all samples. Phospho-peptides were normalized to the corresponding protein abundance value (when available). Additional analysis was done using pathfinder, IPA, and PhosR R packages (Kim et al., 2021; Ulgen et al., 2019).

Supplemental data Tables list all quantified proteins as well as associated TMT reporter ratio to control channels used for quantitative analysis.

### LC-MS/MS with the hybrid metabolomics

Organoids were subjected to an LCMS analysis to detect and quantify known peaks. A metabolite extraction was carried out on each sample based on a previously described method (Pacold et al., 2016). The LC column was a Millipore TMZIC-pHILIC (2.1×150 mm, 5μm) coupled to a Dionex Ultimate 3000TM system and the column oven temperature was set to 25oC for the gradient elution. A flow rate of 100μL/min was used with the 10 mM ammonium carbonate in water (A), pH 9.0, and acetonitrile (B). The gradient profile was 80-20% B (0-30min), 20-80% B (30-31min), 80-80% B (31-42min). Injection volume was set to 2 μL for all analyses (42min total run time per injection). MS analyses were carried out by coupling the LC system to a Thermo Q Exactive HFTM mass spectrometer operating in heated electrospray ionization mode (HESI). Method duration was 30 min with polarity switching data-dependent Top 5method for both positive and negative modes. Spray voltage for both positive and negative modes was 3.5kV and the capillary temperature was set to 320 °C with a sheath gas rate of 35, aux gas of 10, and max spray current of 100 μA. The full MS scan for both polarities utilized 120,000 resolution with an AGC target of 3e^6^ and a maximum IT of 100 ms, and the scan range was from 67-1000 m/z. Tandem MS spectra for both positive and negative modes used a resolution of 15,000, AGC target of 1 e^5^, maximum IT of 50 ms, isolation window of 0.4 m/z, isolation offset of 0.1 m/z, fixed first mass of 50 m/z, and 3-way multiplexed normalized collision energies (nCE) of 10, 35, 80. The minimum AGC target was 1e4 with an intensity threshold of 2e5. All data were acquired in profile mode. The data was analysed using the R Package MetaboAnalyst 4.0 (Chong et al., 2019).

### Glycolysis stress test

Organoids were submitted to the glycolysis stress test in the Agilent Seahorse XF using the glycolysis stress test. Organoids were plated in a laminin-coated Seahorse XFe96 Spheroid Microplate on day 35 and analyzed on day 40 or day 200 after plating. The assays were done in XF base medium supplemented with B27 and N2.

## ACKNOWLEDGMENTS

The authors thank Bill Skarnes, iPSC Neurodegenerative Disease Initiative (iNDI), the Center for Alzheimer’s and Related Dementias (CARD) and the ASAP consortium for the donation of the KOLF2.1J and L166P cell lines. Deans Flow Cytometry CoRE at Icahn School of Medicine Flow Cytometry Core for discussions and technical expertise in sorting, Mount Sinai for flow cytometry training, and the Neuropathology Brain Bank at the Icahn School of Medicine at Mount Sinai for the technical help in sectioning, staining, and imaging spheres. The NYU metabolomics core for help in designing and analyzing the metabolomics experiments. The study is funded by the joint efforts of The Michael J. Fox Foundation for Parkinson’s Research (MJFF) and the Aligning Science Across Parkinson’s (ASAP) initiative. MJFF administers the grant [02663614] on behalf of ASAP and itself., NIH grant R01:0255E131, C.G received a T32: 2T32AG049688-06 grant, L.S. was supported by the Training Program in Stem Cell Biology fellowship from the New York State Department of Health (NYSTEM-C32561GG).

## AUTOR CONTRIBUTIONS

G.M.P, T.D.A, and J.B. design the experiments and wrote the manuscript with input of all authors. G.M.P., E.C., L.S., C.G., R.R., S.S. performed the tissue culture experiments and contributed with data analysis. D.J. contributed with technical expertise and data interpretation for the metabolomics experiments. K.H.N, A.O. contributed with technical expertise and data interpretation for the proteomics experiments. J.F.C., K.W. contributed with design, data analysis and interpretation of human tissue experiments.

## INCLUSION AND DIVERSITY STATEMENT

One or more authors of this paper self-identity as an underrepresented ethnic minority.

---

**Figure S1.**
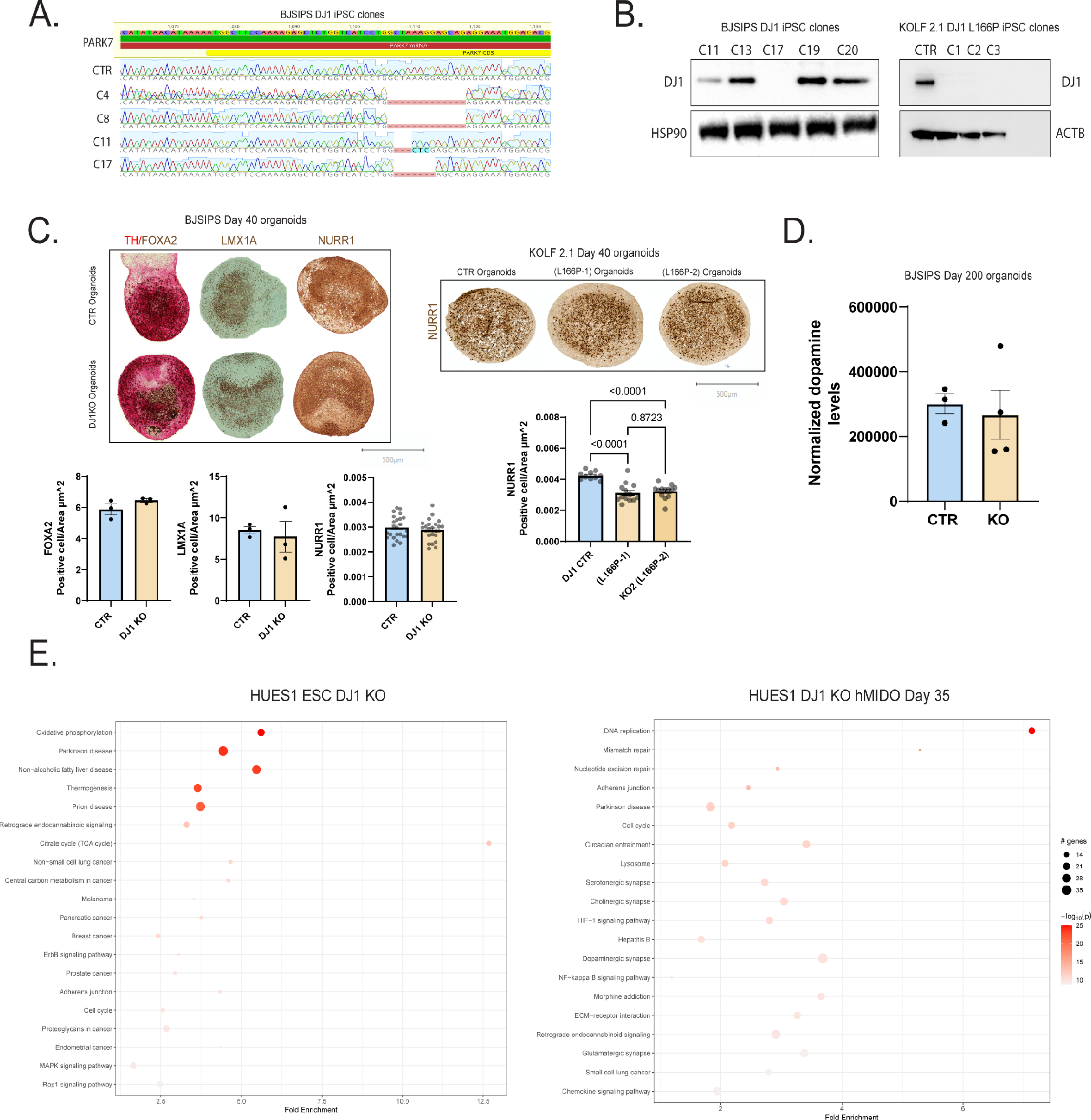
A. Representation of the Sanger sequencing chromatogram for DJ1 CRISPR clones in BJ-SIPS iPSC line. B. KO confirmation immunoblot for DJ1 CRISPR clones in BJ-SIPS and KOLF 2.1J iPSC lines. C. Midbrain organoids staining and quantification of TH/FOXA2 and LMX1A markers at day 20 and NURR1 at day 40 positive cells. D. Mass spectrometry dopamine quantification of midbrain organoids at day 200. E. Pathway enrichment analysis in DJ1 KO generated from HUES1 iPSC and day 35 midbrain organoids. All data are represented in mean ± S.E.M, data points are individual organoids, and the p-value was reported on the graph highlighted comparison.

**Figure S2.**
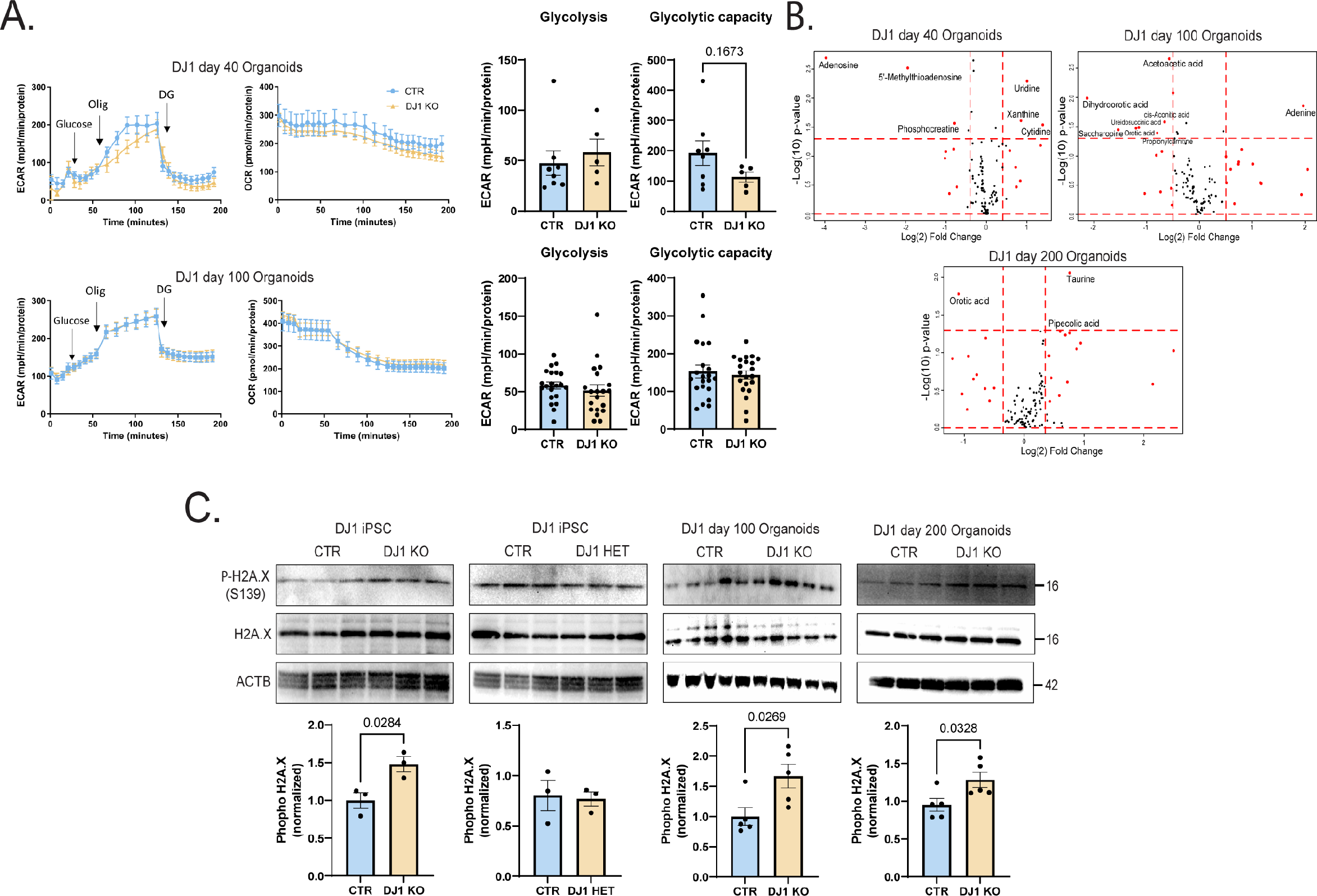
A. Glycolysis stress seahorse assay in day 40 and day 100 midbrain organoids showing ECAR and OCR levels pre-glucose and after glucose, oligomycin and DG treatments. B. Volcano plot representation of metabolomics panel for energy-related metabolites in day 40, 100, and 200 midbrain organoids. C. Immunoblots for native H2AX, phospho-H2AX (S139), and actin (ACTB) loading control for CTR and DJ1 KO iPSC and day 100 and 200 midbrain organoids. All data are represented in mean ± S.E.M, data points from seahorse assay are individual organoids, and data points for Wb are individual well differentiations. p-value was reported on the graph highlighted comparison.

**Figure S3.**
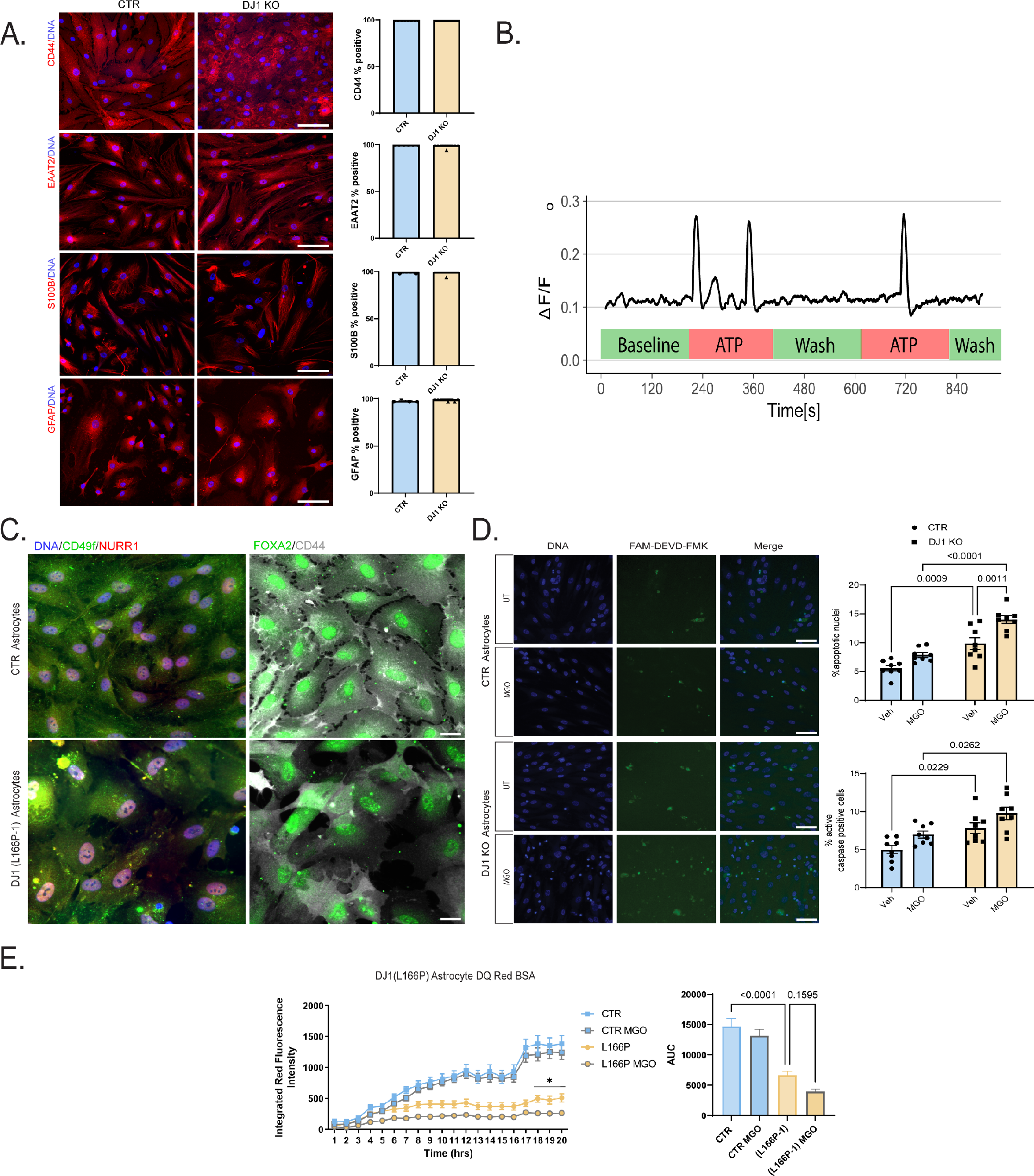
A. Immunostaining panel of CD44, EAAT2, S100B, and GFAP counterstained with DAPI in midbrain astrocytes and % of positive cells. B. Astrocyte GCAMP8s calcium imaging δF/F trace before and after ATP stimulation. C. Immunostaining panel of midbrain markers NUUR1 and FOXA2 co-stained with CD44. D. Poli-caspase reporter peptide FAM-DEVD-FMK staining in MGO treated or untreated CTR or DJ1 KO astrocytes counterstained with DAPI. Quantitation of the standing showing percentage apoptotic nuclei and caspase positive cells. E. DQBSA proteolysis live imaging assay of vehicle or MGO treated astrocytes of CTR or DJ1 L166P genotypes. Scale bars, 100 μm for A.; 20 μm for C; 50 μm for D. All data are represented in mean ± S.E.M, data points are individual organoids, and the p-value was reported on the graph highlighted comparison.

**Figure S4.**
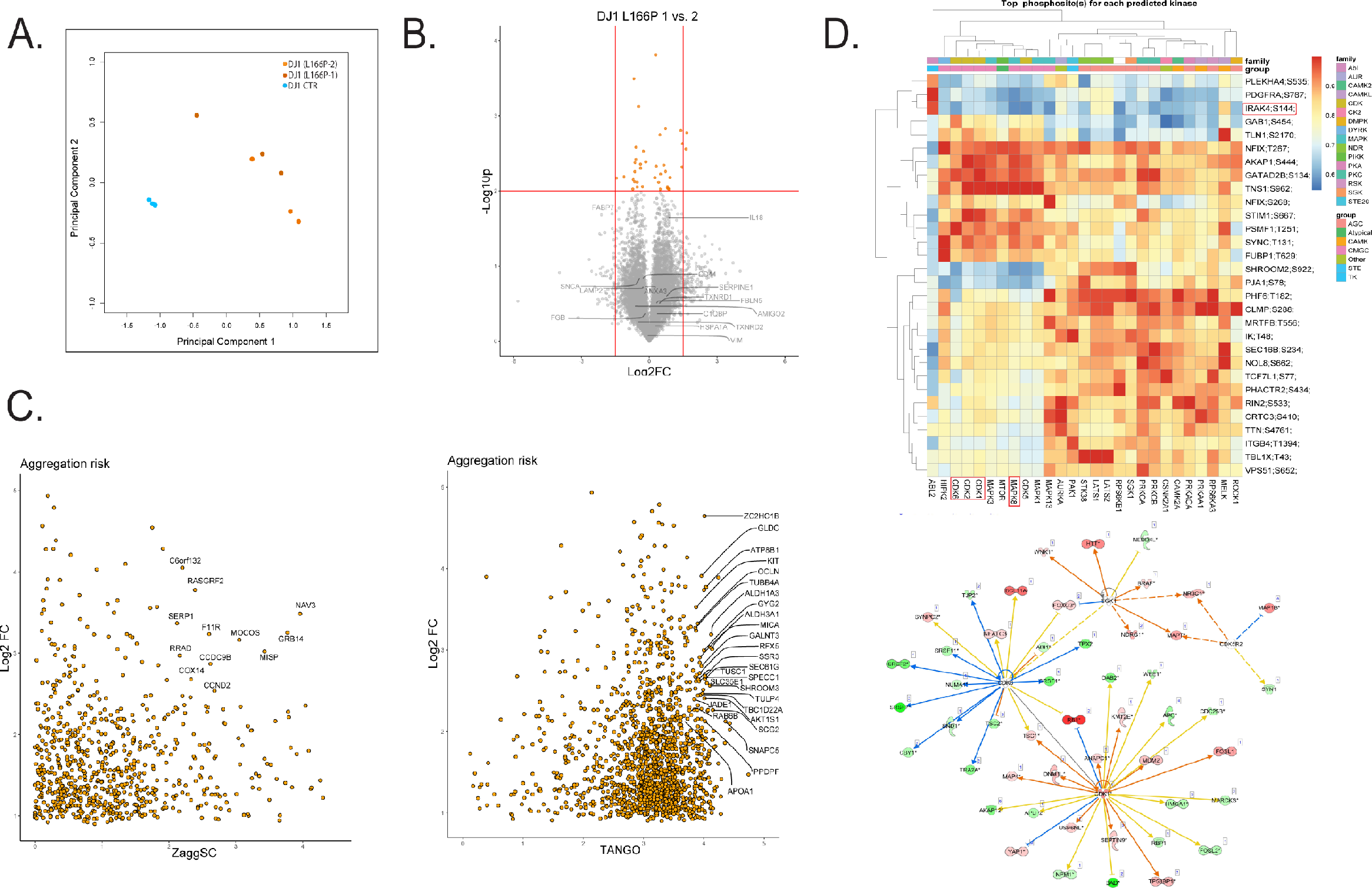
A. PCA analysis of the TMT-labelled proteomics in KOLF 2.1J CTR, DJ1 L166P KO1, and KO2 midbrain astrocytes. B. Volcano plot comparing the two DJ1 L166P clones highlighting selected proteins. C. Log2FC DJ1 L166P KO1 proteomics integration with aggregation risk scores for the human proteome ZaggSC and D. TANGO score showing selected proteins. D. Clustering plot of phospho-proteomics kinase activity prediction showing the top phosphosites and IPA Kinase/phosphosite network plots.

**Figure S5.**
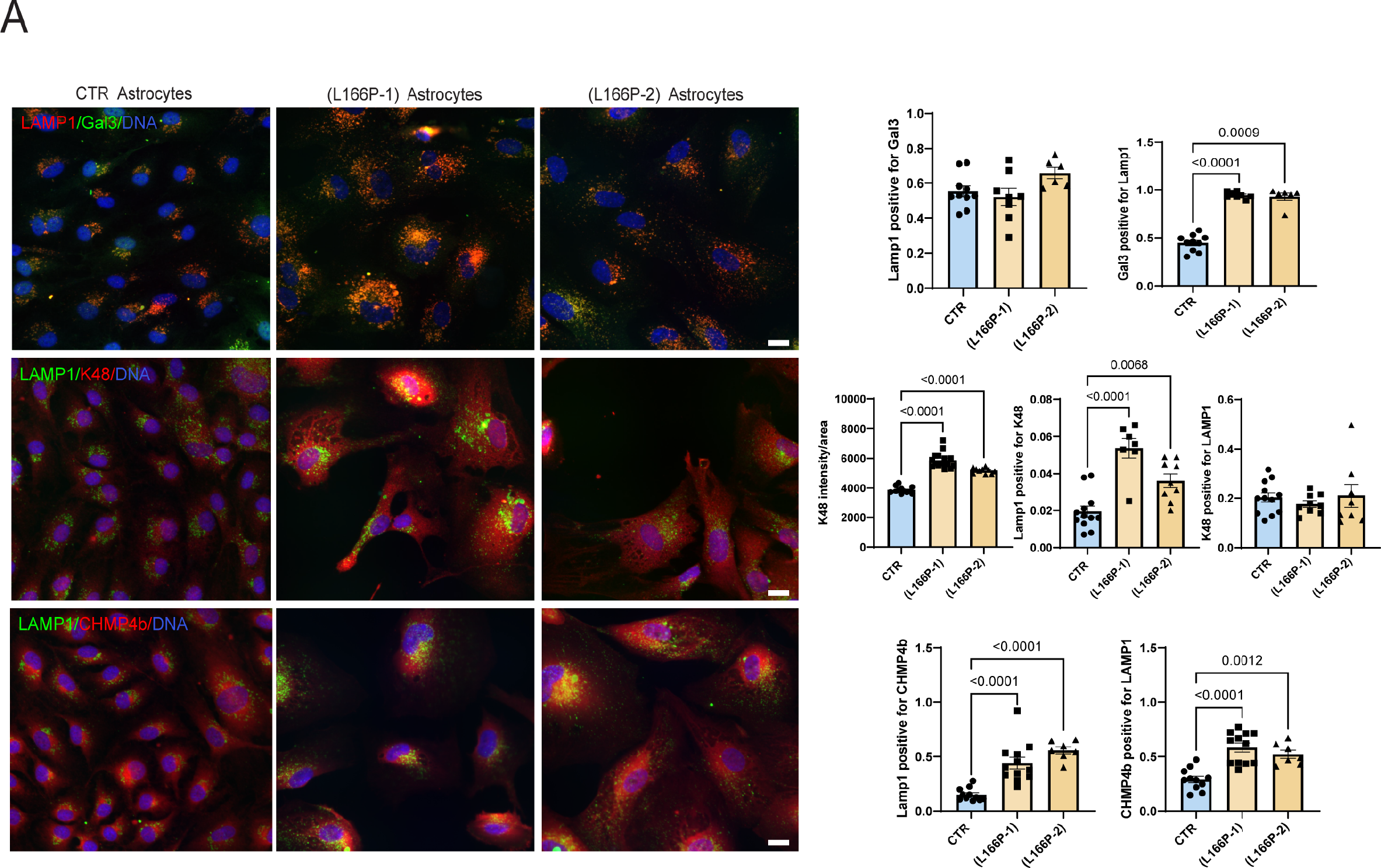
A. Immunostaining panel showing Gal3 co-stained with LAMP1 in KOLF 2.1J CTR, DJ1 L166P KO1, and KO2 midbrain astrocytes. Co-localization analysis of Gal3 co-stained with LAMP1. Immunostaining panel showing K48 Ub chain co-stained with LAMP1 in KOLF 2.1J CTR, DJ1 L166P KO1, and KO2 midbrain astrocytes. Co-localization analysis of K48 Ub chain co-stained with LAMP1. Immunostaining panel showing CHMP4b co-stained with LAMP1 in KOLF 2.1J CTR, DJ1 L166P KO1, and KO2 midbrain astrocytes. Co-localization analysis of CHMP4b co-stained with LAMP1. Scale bars, 15 μm. All data are represented in mean ± S.E.M, data points are individual wells, and the p-value was reported on the graph highlighted comparison.

