## Supplementary material for "Disruption of lysosomal proteolysis in astrocytes facilitates midbrain proteostasis failure in an early-onset PD model": Table 2

| REAGENT or RESOURCE | SOURCE | IDENTIFIER |
| --- | --- | --- |
| Antibodies |  |  |
| FOXA2 | Abcam | Cat#Ab108422; RRID: AB_11157157 |
| GAPDH | Abcam | Cat#Ab9485; RRID: AB_307275 |
| GFAP | Millipore | Cat#MAB360; RRID: AB_11212597 |
| TH | Millipore | Cat#MAB318; RRID: AB_2201528 |
| TH | ABCAM | Cat#ab112;RRID: N/A |
| FOXA2 | ABCAM | Cat#ab40874;RRID: N/A |
| PARK7/DJ1 | ABCAM | Cat#ab169520;RRID: N/A |
| LMX1A | SIGMA | Cat#HPA030088;RRID: N/A |
| FOXA2 | ABCAM | Cat#ab60721;RRID: N/A |
| TH | Millipore | Cat#MAB318; RRID:AB_2201528 |
| alpha Synuclein | ABCAM | Cat#ab138501;RRID:AB_2537217 |
| alpha Synuclein (Phosphorylated S129) | ABCAM | Cat#ab168381;RRID:AB_2728613 |
| alpha Synuclein (Filament) | ABCAM | Cat#ab209538;RRID:AB_2714215 |
| EAAT2 | ABCAM | Cat#ab41621;RRID: N/A |
| LC3A/B | Cell Signaling | Cat#4108S;RRID: N/A |
| RAGE | ABCAM | Cat#ab37647;RRID:AB_777613 |
| Histone H2A.X | Millipore | Cat#07-627; RRID:AB_2233033 |
| CD49f | Biolegend | Cat#313602;RRID:AB_345296 |
| Methylglyoxal (MGO) | Cell Biolabs | Cat#STA-011,RRID: N/A |
| N-epsilon-CML | Cell Biolabs | Cat#STA-014;RRID: N/A |
| S100 - Beta | ABCAM | Cat#ab52642;RRID:AB_882426 |
| LAMP1 | ABCAM | Cat#ab25630;RRID:AB_470708 |
| LC3B | Cell Signaling | 3868S;RRID: N/A |
| NR4A2 (NURR1) | SIGMA | Cat#N6413;RRID:AB_1841046 |
| CHMP4B | ptglab | Cat#13683-1-AP;RRID: N/A |
| Galectin-3 (Mac-2) | Biolegend | Cat#125401;RRID:AB_1134237 |
| CD44 | ABCAM | Cat#ab157107;RRID:AB_2847859 |
| Ubiquitin (Lys48-Specific), clone Apu2 | Millipore | Cat#05-1307;RRID:AB_1587578 |
| Phospho-Histone H2A.X (Ser139), clone JBW301 | Millipore | Cat#05-636;RRID:AB_309864 |
| oxDJ-1 (Cys106), clone M149 | Millipore | Cat#MABN1773;RRID: N/A |
| β-Actin (C4) HRP | Santa Cruz | Cat#sc-47778;RRID:AB_2714189 |
| P62 | Progen | Cat#GP62-C;RRID:AB_2687531 |
| GBA | Abnova | Cat#H00002629-M01;RRID:AB_464151 |
| EAAT2 | ABCAM | Cat#ab41621;RRID:AB_941782 |
| TUBB3 | Abcam | Cat#Ab78078; RRID: AB_2256751 |
| Chemicals, peptides, and recombinant proteins |  |  |
| Accutase® Cell Detachment Solution | Innovative Cell Technologies | Cat#AT104; RRID: AB_2869384 |
| B-27™ Supplement (50×), Minus Vitamin A | Gibco | Cat#12587010; RRID: N/A |
| Bovine Albumin Fraction V (7.5% Solution) | Gibco | Cat#15260037; RRID: N/A |
| CHIR99021 | Tocris | Cat#4423; RRID: N/A |
| DAPT | Cayman Chemical | Cat#13197; RRID: N/A |
| DB-cAMP/Dibutyryl-cAMP | Biolog | Cat#D 009; RRID: N/A |
| DirectPCR Lysis Reagent (Cell) | Viagen Biotech | Cat#302-C; RRID: N/A |
| DMEM/F-12, GlutaMAX™ Supplement | Gibco | Cat#10565018; RRID: N/A |
| Astrocyte Medium | Sciencell | Cat#1801; RRID: N/A |
| DMEM, High Glucose | Gibco | Cat#11965092; RRID: N/A |
| Geltrex™ LDEV-Free, hESC-Qualified, Reduced Growth Factor Basement Membrane Matrix | Gibco | Cat#A1413302; RRID: N/A |
| Hoechst 33342, Trihydrochloride, Trihydrate - 10 mg/mL Solution in Water | Invitrogen | Cat#H3570; RRID: N/A |
| Laminin Mouse Protein, Natural | Gibco | Cat#23017015; RRID: N/A |
| L-Ascorbic Acid (White Crystalline Powder) | Fisher Scientific | Cat#BP351-500; RRID: N/A |
| LDN193189 | Stemgent | Cat#04-0074; RRID: N/A |
| Leibovitz's L-15 Medium | Gibco | Cat#11415064; RRID: N/A |
| N-2 Supplement (100×) | Gibco | Cat#17502048; RRID: N/A |
| Opti-MEM™ I Reduced Serum Medium | Gibco | Cat#31985062; RRID: N/A |
| PBS, pH 7.4 (-CaCl2, -MgCl2) | Gibco | Cat#10010023; RRID: N/A |
| Penicillin-Streptomycin (10,000 U/mL) | Gibco | Cat#15140122; RRID: N/A |
| Poly-L-Ornithine Hydrobromide | Sigma-Aldrich | Cat#P3655; RRID: N/A |
| Proteinase K Solution (20 mg/mL) | Viagen Biotech | Cat#501-PK; RRID: N/A |
| Purmorphamine | STEMCELL Technologies | Cat#72204; RRID: N/A |
| Puromycin Dihydrochloride | Gibco | Cat#A1113803; RRID: N/A |
| Recombinant Human BDNF Protein | R&D Systems | Cat#248-BD; RRID: N/A |
| Recombinant Human CNTF Protein | Peprotech | Cat#450-13;RRID: N/A |
| Recombinant Human GDNF Protein | R&D Systems | Cat#212-GD; RRID: N/A |
| SAG | Cayman Chemical | Cat#11914; RRID: N/A |
| SB431542 | Stemgent | Cat#04-0010-10; RRID: N/A |
| StemFlex™ Medium | Gibco | Cat#A3349401; RRID: N/A |
| Thiazovivin | Selleck Chemicals | Cat#S1459; RRID: N/A |
| Trypan Blue Solution, 0.4% | Gibco | Cat#15250061; RRID: N/A |
| Aminoguanidine hydrochloride | Sigma | Cat#1937-19-5; RRID: N/A |
| Bafilomycin A1 from Streptomyces griseus | Sigma | Cat#B1793; RRID: N/A |
| (R)-MG-132 | Selleck Chemicals | Cat#S2619; RRID: N/A |
| Non-essential Amino Acids (NEAA) | Gibco | Cat#11140050;RRID: N/A |
| GlutaMax Supplement | Gibco | Cat#35050061;RRID: N/A |
| Triton X-100 | Sigma | Cat# T9284;RRID: N/A |
| RIPA buffer | Sigma | Cat# R0278;RRID: N/A |
| DQ™ Red BSA | ThermoFisher | Cat#D12051;RRID: N/A |
| Y-27632 Dihydrochloride | Tocris | Cat#1254; RRID: N/A |
| Critical commercial assays |  |  |
| Seahorse XF Glycolysis Stress Test Kit | Agilent | Cat#103020-100;RRID: N/A |
| Image-iT™ LIVE Red and Green Caspase Apoptosis Detection Kits for microscopy | ThermoFisher | Cat#I35101;RRID: N/A |
| Lipofectamine™ Stem Transfection Reagent | Invitrogen | Cat#STEM00003; RRID: N/A |
| PROTEOSTAT | Enzo | Cat#ENZ-51023;RRID: N/A |
| P3 Primary Cell 4D-Nucleofector™ X Kit S | Lonza | Cat#V4XP-3032; RRID: N/A |
| Experimental models: Cell lines |  |  |
| Human: BJ SiPS-D induced pluripotent stem cells | Harvard University | Cat#BJ SiPS-D: RRID: CVCL_X741 |
| KOLF2.1J | Jackson Laboratory | RRID: N/A |
| KOLF2.1J DJ1 L166P 1 | Jackson Laboratory | RRID: N/A |
| KOLF2.1J DJ1 L166P 2 | Jackson Laboratory | RRID: N/A |
| Human: BJ SiPS-D induced pluripotent stem cells Dj1 HET #11 | This paper | RRID: N/A |
| Human: BJ SiPS-D induced pluripotent stem cells Dj1 KO #17 | This paper | RRID: N/A |
| Oligonucleotides |  |  |
| PARK7 _L166P F      ATGCATACCCGCCTCCATTACGTTG | This paper | RRID: N/A |
| PARK7 _L166P R      ATAAGCAGAGAAAATCACAAGCCTC | This paper | RRID: N/A |
| PARK7 _L166P seq        AATGGATTCCTAACGGCCTG | This paper | RRID: N/A |
| PARK7 R ATGGCTAAAAATCGATGTGG | This paper | RRID: N/A |
| PARK7 F TGGGGTATCTCAGGGTTGCA | This paper | RRID: N/A |
| Recombinant DNA |  |  |
| Plasmid: pTAHR TH-p2a-TD:Tomato (floxed selection Puro) | Ahfeldt et al., 2020; Addgene | Cat#135814; RRID: N/A |
| CRISPR 1: pTACR TH1-p2aGFP | Ahfeldt et al., 2020; Addgene | Cat#135815; RRID: N/A |
| CRISPR 2: pTACR TH2-p2aGFP | Ahfeldt et al., 2020; Addgene | Cat#135816; RRID: N/A |
| Plasmid: pCAG-Cre:GFP | Matsuda and Cepko, 2007; Addgene | Cat#13776; RRID: Addgene_13776 |
| Software and algorithms |  |  |
| CellProfiler | https://cellprofiler.org/ | RRID:SCR_007358 |
| Rstudio | https://www.rstudio.com/ | RRID:SCR_000432 |
| Ingenuity Pathway Analysis (IPA) | Qiagen | RRID:SCR_008653 |
| QuPath | https://QuPath.github.io/ | RRID:SCR_018257 |
| Pathfinder | https://github.com/egeulgen/pathfindR | Ulgen et al., 2019 |
| PhosR R packages | https://github.com/PYangLab/PhosR | Kim et al., 2021 |
| MetaboAnalyst 4.0 | https://www.metaboanalyst.ca | RRID:SCR_015539 |
| Fiji | https://fiji.sc/ | Chong et al., 2019; RRID:SCR_002285 |
| Other |  |  |
| 4D-Nucleofector™ Core Unit | Lonza | Cat#AAF-1002B; RRID: N/A |
| Cell Culture Microplate, 96-well, PS, F-bottom (Chimney Well), μClear®, Black, CELLCOAT®, Poly-L-Lysine, Lid with Condensation Rings, 5 pcs./bag | Greiner | Cat#655936; RRID: N/A |
| Corning® 125 mL Disposable Spinner Flask with 70 mm Top Cap and 2 Angled Sidearms, Sterile | Corning | Cat#3152; RRID: N/A |
| Corning® Costar® Ultra-Low Attachment Multiple Well Plate | MilliporeSigma | Cat#CLS3471; RRID: N/A |
| Countess™ Cell Counting Chamber Slides | Invitrogen | Cat#C10228; RRID: SCR_019815 |
| Countess™ II FL Automated Cell Counter | Invitrogen | Cat#AMQAF1000; RRID: N/A |
| Falcon® Round-Bottom Tubes with Cell Strainer Cap, 5 mL | STEMCELL Technologies | Cat#100-0087; RRID: N/A |
| Fisherbrand™ Disposable Borosilicate Glass Pasteur Pipets | Fisher Scientific | Cat#13-678-20D; RRID: N/A |
| Fisherbrand™ Sterile Cell Strainers (70 μm) | Fisher Scientific | Cat#22-363-548; RRID: N/A |
| pluriStrainer® 300 μm, 25 pcs. – Sterile (Cell Strainer) | pluriSelect | Cat#43-50300-03; RRID: N/A |
| pluriStrainer® 500 μm, 25 pcs. – Sterile (Cell Strainer) | pluriSelect | Cat#43-50500-03; RRID: N/A |
| Stirrers, Magnetic, Nine-position, Dura-Mag | Chemglass Life Sciences | Cat#CLS-4100-09; RRID: N/A |
| Variable Speed 2D Rocker | USA Scientific | Cat#2527-2000; RRID: N/A |
